# Activation of neuronal FLT3 promotes exaggerated sensorial and emotional pain-related behaviors facilitating the transition from acute to chronic pain

**DOI:** 10.1101/2021.11.23.469557

**Authors:** Adrien Tassou, Maxime Thouaye, Damien Gilabert, Antoine Jouvenel, Jean-Philippe Leyris, Corinne Sonrier, Lucie Diouloufet, Ilana Mechaly, Sylvie Mallié, Myriam Chentouf, Madeline Neiveyans, Martine Pugnière, Pierre Martineau, Bruno Robert, Xavier Capdevila, Jean Valmier, Cyril Rivat

## Abstract

Acute pain events have been associated with persistent pain sensitization of nociceptive pathways increasing the risk of transition from acute to chronic pain. However, it is unclear whether injury-induced persistent pain sensitization can promote long-term mood disorders. The receptor tyrosine kinase FLT3 is causally required for pain chronification after peripheral nerve injury, questioning its role in the development of pain-induced mood alterations. Here, we evaluated the emotional and sensorial components of pain after a single (SI) or double paw incision (DI). We then investigated the role of FLT3 either by inhibition using transgenic knock-out mice and functional antibodies or by activation with FLT3 ligand (FL) administrations.

DI mice showed significant anxiodepressive-like and spontaneous pain behaviors as opposed to SI mice. DI also promoted and extended mechanical pain hypersensitivity compared to SI. This emotional and sensorial pain exaggeration correlated with spinal changes especially by increased microglia activation after DI versus SI. Intrathecal minocycline, a microglial inhibitor, specifically reversed DI induced-mechanical hypersensitivity in males. Repeated treatment with the microglia proliferation inhibitor GW2580 not only eliminated the exaggerated pain hypersensitivity produced by DI but also prevented anxiodepressive-related behaviors in DI animals. Finally, FL injections in naive animals provoked mechanical allodynia and anxiodepressive-like disorders concomitant with a strong microglial activation while *Flt3* silencing in a genetic mouse line or FLT3 blocking via functional antibodies, blunted the development of persistent pain and depression after DI. Altogether our results show that the repetition of peripheral lesions facilitate not only exaggerated nociceptive behaviors but also induced anxiodepressive disorders supported by spinal central changes. The inhibition of FLT3 could thus become a promising therapy in the management of pain sensitization and related mood alterations.

## Introduction

Understanding the transition from acute to chronic pain remains an important challenge to better manage pain in patients [1]. This is specifically true in the context of surgery where chronic postsurgical pain (CPSP) is significantly debilitating, under-evaluated and affecting several millions of people each year in the world [2–5]. Pre-existing pain is a major risk factor as it profoundly modifies patients internal homeostasis altering the degree of pain sensitization [6–8]. Hence, sensitization after sensorial trauma appears to be a critical event in the development of chronic pain by amplifying signaling in previously affected nociceptive pathways [9]. Animal models called “hyperalgesic priming” are classically used to study the neuroplasticity underlying persistent pain sensitization [10– 18]. These models consist in performing an initial nociceptive stimulation that induces a prolonged period of susceptibility to exaggerated pain behavior after a subsequent stimulation. This phenomenon has been named latent pain sensitization [19] and its mechanisms have been further studied [20–22]. Furthermore, the clinical observation that chronic pain incidence after injury is largely related to preinjury pain status [6] suggests that the maintenance of latent sensitization may facilitate the transition from acute to persistent pain. Mood disorders such as depression and anxiety are frequently observed in patients suffering from chronic pain. It has been reported that at least 50% of chronic pain patients experience a major depressive disorder [23], making depression and chronic pain together a major comorbidity [24–26]. Similarly, persistence of mood disorders is a maladaptive process maintained by neuropathological mechanisms that can aggravate the sensory abnormalities of chronic pain [27, 28]. Although this comorbidity is clinically well established, the underlying pathophysiological mechanisms remain poorly understood. It has been proposed that depression is a consequence of the presence of chronic pain [29, 30] and that peripheral nociceptive inputs may act as a trigger for central mechanisms that leave predisposed individuals at increased risk of anxiodepressive disorders. Thus, we hypothesize that peripheral nociceptive inputs may generate sustained neurochemical alterations (latent sensitization) leading to affective disorders after subsequent re-exposure to the same nociceptive stimulus as shown with exaggerated pain hypersensitivity in hyperalgesic priming models. To date, the affective dimension has never been fully studied in animal models of hyperalgesic priming and this incomplete characterization has impeded the understanding of the role of peripheral nociceptive inputs in the development of affective disorders. Consequently, we characterized a model of repeated hindpaw injuries where the animals undergo two hind paw surgery 7 days apart, by studying both sensory and affective alterations.

Recently, we reported the expression of the fms-like tyrosine kinase 3 receptor (FLT3) in the peripheral nervous system [31] and its implication in the development and maintenance of chronic neuropathic pain *via* its ligand, the cytokine FL (FLT3 Ligand). Originally identified as part of the hematopoietic system, we showed that FLT3, expressed in DRG neurons, induces long-term molecular modifications, leading to neuronal hyperexcitability giving rise to neuropathic pain symptoms. In contrast, the genetic and pharmacological inhibition of FLT3 prevents, but also reverses, hyperexcitability and pain-related behaviors associated with nerve injury. These results strongly highlight the key role of FLT3 in the development of long-term neuropathic pain sensitization. Although its role in the sensorial component of neuropathic pain is now established, we are still lacking evidence about the extent to which peripheral FLT3 can be involved in the central consequences of latent sensitization, especially in the affective dimension of pain. Hence, in a second objective, we revealed a FLT3 dependent peripheral sensitization that triggers both exaggerated nociceptive and affective disorders after repeated surgical injury.

## Methods and materials Animals

Experiments were performed in C57BL/6 naive mice (Janvier, France), mice carrying a homozygous deletion of *Flt3* (*Flt3*^-/-^ mice) [32] or *CX3CR1*^*EGFP*^ mice and their littermates (WT) weighing 25-30 g. All the procedures were approved by the French Ministry of Research (authorization #1006). Animals were maintained in a climate-controlled room on a 12 h light/dark cycle and allowed access to food and water *ad libitum*. Male and female mice were first considered separately in behavioral procedures. Both sexes showed mechanical hypersensitivity of same intensity after intrathecal FL injection and after surgery and were similarly affected by *Flt3* deletion (ANOVA followed by Bonferroni’s test, n = 8 for both sexes and genotypes for each experiment). With some exceptions shown in the results, experiments were performed mainly on male mice.

### Surgeries

Paw incision models: Mice C57Bl6/J were anesthetized under isoflurane (3% vol/vol). For the single incision (SI) model [33], a 0.7cm incision was applied with a number 11 blade on the skin and fascia of the left hindpaw plantar surface. Subcutaneous plantaris muscle was then isolated, exposed and longitudinally incised. After hemostasis, the wound was sutured with two 6.0 absorbable sutures and the mice were finally placed in recovery cages. For the double incision (DI) model [34], the same procedure was repeated on the opposite hindpaw 1 week after the first incision. Control animals underwent a Sham procedure, which consisted in isoflurane (3% vol/vol) anesthesia alone.

Chronic constriction injury (CCI) model: CCI of the sciatic nerve was performed as described previously [35] and adapted for mice [36].

### Injections

FL injections: To study FL induced microgliosis, human FL (50ng/5μl) produced as previously described (6) or vehicle was intrathecally injected daily for 3 days, spinal cords were then collected for immunohistochemistry. To know whether FL can induce anxiodepressive-like disorders, mice underwent FL or vehicle intrathecal injection (50ng/5μl) every 3 days during 15 days and were subjected to anxiodepression testing 24 hours after the last injection. Curative minocycline was intrathecally injected 3 days post-incision or sham at a dose of 300μg/5μl/mouse. Preventive GW2580 was intrathecally injected during surgery or sham at a dose of 1μg/5μl/mouse. Preventive administration of mAbA3 consisted in a single intraperitoneal injection (200 μg in 200 μl) during the first incision. Repeated injections consisted in 1 injection every 2 days during the first incision phase for a total of 4 injections for the evaluation of anxiodepressive-like behaviors 2 weeks post-injury.

### Production of human recombinant FL (rh-FLT3-L)

See Supplementary Methods and Materials for details.

### Anti-FLT3 antibody development and production

See Supplementary Methods and Materials for details.

### Affinity measurements of Anti-FLT3 antibody for human and murine FLT3

The surface plasmon resonance (SPR) experiments were performed on a T200 apparatus at 25°C on CM5S dextran sensor chip in HBS-EP+ buffer (20mM Hepes Buffer pH7.4 containing 150mM of NaCl and 0.05% P20 surfactant). Recombinant human and murine FLT3 hFc Chimeric (R&Dsystems) were captured on anti-human Fc using Ab human Capture Kit (Cytiva) according to the manufacturer’s instructions (Cytiva) and increasing concentrations (from 0.75 to 100mM) of anti-FLT3 were injected at 50μl/min. A pulse of 3M MgCl2 was used as regenerant between cycles. The kinetic constants were evaluated from the sensorgrams after double-blank subtraction with T200 Evaluation software (Cytiva) using a bivalent fitting model.

### Behavioral testing

See Supplementary Methods and Materials for details.

### Retrograde labeling

10μl of 1% Fluorogold was applied on the open wound incision during surgery. DRGs were then dissected 7 days after in PBS and post-fixed for 15 minutes in 4% paraformaldehyde (PFA). Tissues were rinsed twice in PBS before immersion overnight in 25% sucrose/PBS at 4°C and were finally frozen. Frozen DRGs were then subjected to cryosection and imaging. Images were collected using a Carl Zeiss LSM 700 confocal microscope and were processed with ZEN software.

### Immunohistochemistry

Immunostainings were performed with the following antibodies (see table 1): ATF3 (Santa Cruz), CSF1 (R&D), Iba1 (Wako), GFAP (abcam), CGRP (abcam), NeuN (Millipore), BrdU (Abcam), and fluorophore coupled secondary antibodies (Alexa Fluor 488, 555, 594, 647) (Invitrogen). Images were collected with a Carl Zeiss LSM 700 microscope or a Zeiss Axioscan slides scanner and were processed with Fiji/ImageJ (NIH). Corresponding images (e.g. ipsilateral vs. contralateral; FL vs vehicle; wt vs. mutant) were processed in an identical manner. For BrdU detection, slides were incubated in 10 mM sodium citrate buffer (pH 6.0) at 95°C for 15 min, then at room temperature (RT) for 15 min and subjected to immunostaining.

See Supplementary Methods and Materials for more details.

### RNA isolation and real time polymerase chain reaction

Dorsal spinal cords were dissected, ipsilateral and contralateral parts were separated and tissues were stored at -80°C until RNA extraction with the TRI reagent/ chloroform technique following the protocol of the manufacturer (sigma). After extraction, samples were treated with DNAse (M610A, promega) and reverse transcribed. Q-PCR reactions were made in 96 well plates with a final volume of 10μl composed of 3μl cDNA (final dilution 1 : 90), 0.5μM of forward and reverse primers and 5μl of SybrGreen I master mix (Roche life Sciences). Measures were realized with a LightCycler 480 (Roche). Results were normalized with the geometric mean of 2 housekeeping genes, *Ddx48* and *Ywaz*, defined beforehand with a screening of 10 housekeeping genes on our samples. Relative quantification of PCR products was realized with the delta-CT method [37].

### In situ hybridization

Antisense digoxigenin (DIG)-labeled RNA probe was generated from a mouse Flt3 cDNA clone (IRAMp995N1310Q, GenomeCUBE Source Bioscience) in a 20 μl reaction containing 1 μg of linearized plasmid (digested with NheI) using the DIG RNA labeling mix (Roche Diagnostics) and Sp6 RNA polymerases (Promega) following the manufacturer’s instructions. DIG-labeled RNA probe was purified on MicroSpin G50 columns (GE Healthcare). Naive or DI-injured WT and CX3CR1^EGFP^ mice were euthanized by Euthasol vet injection (25μl). Spinal cords were dissected in phosphate-buffered saline (PBS) and fixed for 30 min in 4% PFA at room temperature. Tissues were rinsed twice in PBS before immersion overnight in 25% sucrose/PBS at 4°C. In situ hybridization was performed with standard procedures on transverse sections of 14 μm as described [38]. Image acquisition was done using an Axioscan slides scanner ZEISS.

### HTRF binding assay

Tag-lite binding assays were performed 24 h after transfection on fresh cells in 96-well plates. Cells were incubated with 0.5 nM Red-FL in the presence of increasing concentrations of anti-FLT3 mAb to be tested and incubated for 1h at Room temperature prior to the addition of Red-FL. Plates were then incubated at room temperature overnight before signal detection.

### HTRF autophosphorylation assay

RS4-11 cells were used to carry out this assay in suspension in 384-well small volume plates to determine the degree of phosphorylation of FLT3 receptors. In the plates containing RS4-11 cells, anti-FLT3 mAb to be tested were added and incubated for 90min at room temperature prior to the addition of rh-FL at 10 or 1 nM for 10 min at room temperature. After incubation, cells were lysed and a mix of anti-FLT3 and anti-TYR-969 FLT3 antibodies labeled with Lumi4-Tb and d2 respectively were added to the cell lysate. Plates were then incubated at room temperature for 2h before signal detection. Autophosphorylation measured in the absence of FL represents FLT3 constitutive activity whereas that measured in the presence of FL represents FL-induced activity.

### Statistical analysis

Data were analyzed with graphpad prism 6. Area Under Curve (AUC) was calculated based on the trapezoidal rule to have the volume of area above or under the baseline across different time points until full recovery. Data are expressed as the mean ± S.E.M. All sample sizes were chosen based on our previous studies except for animal studies for which sample size has been estimated via a power analysis using the G-power software. The power of all target values was 80% with an alpha level of 0.05 to detect a difference of 50%. Statistical significance was determined by analysis of variance (ANOVA one-way or two-way for repeated measures, over time). In all experiments in which a significant result was obtained, the test was followed by Holms-Sidak post-hoc test for multiple comparisons, as appropriate. In case of two experimental groups, unpaired two-tailed t-test was applied. For cell counting, and area quantification, statistical analyses were performed using Mann-Whitney test. The applied statistical tests are specified in each figure legend.

## Results

### Characterization of sensorial and emotional-related behaviors after repeated hindpaw injuries

SI (Single Incision, one incision on the left hindpaw) [33] and DI (Double Incision, first incision on the left hindpaw and second one on the right hindpaw) models were compared and tested at three different time points after surgery: the following days after surgery (1^st^ week post-incision), just after nociceptive recovery (2^nd^ week post-incision) and 1 month after incision. Because these behaviors require motor functions, motor coordination was systematically assessed and found unchanged (Fig. S1A). The first week after surgery, SI and DI mice responded to the NSF with an increased latency to eat compared to Sham mice while only DI mice responded to ST and FST with a decreased grooming behavior and immobility, respectively (Fig. S1B-D). The second week after surgery, we found no difference between Sham mice and SI mice in all the tests while DI mice showed sustained anxiodepressive-like behaviors (Fig. 1A-C). One month after surgery, all the experimental groups returned to Sham values (Fig. S1E-G). These results are resumed in Fig. 1D. Decreased neurogenesis at the level of the dentate gyrus has been associated with depression in human [39] and animal [40, 41] studies. We therefore hypothesized that DI-induced anxiodepressive-like behaviors could decrease the rate of newborn neurons in this region as evaluated with Brdu staining. Brdu stained neurons (Brdu/NeuN colocalization) were substantially decreased in DI mice 7 days after the surgery compared to SI and Sham mice (Fig. 1E-F) supporting the development of mood disorder in DI mice.

**Figure 1:**
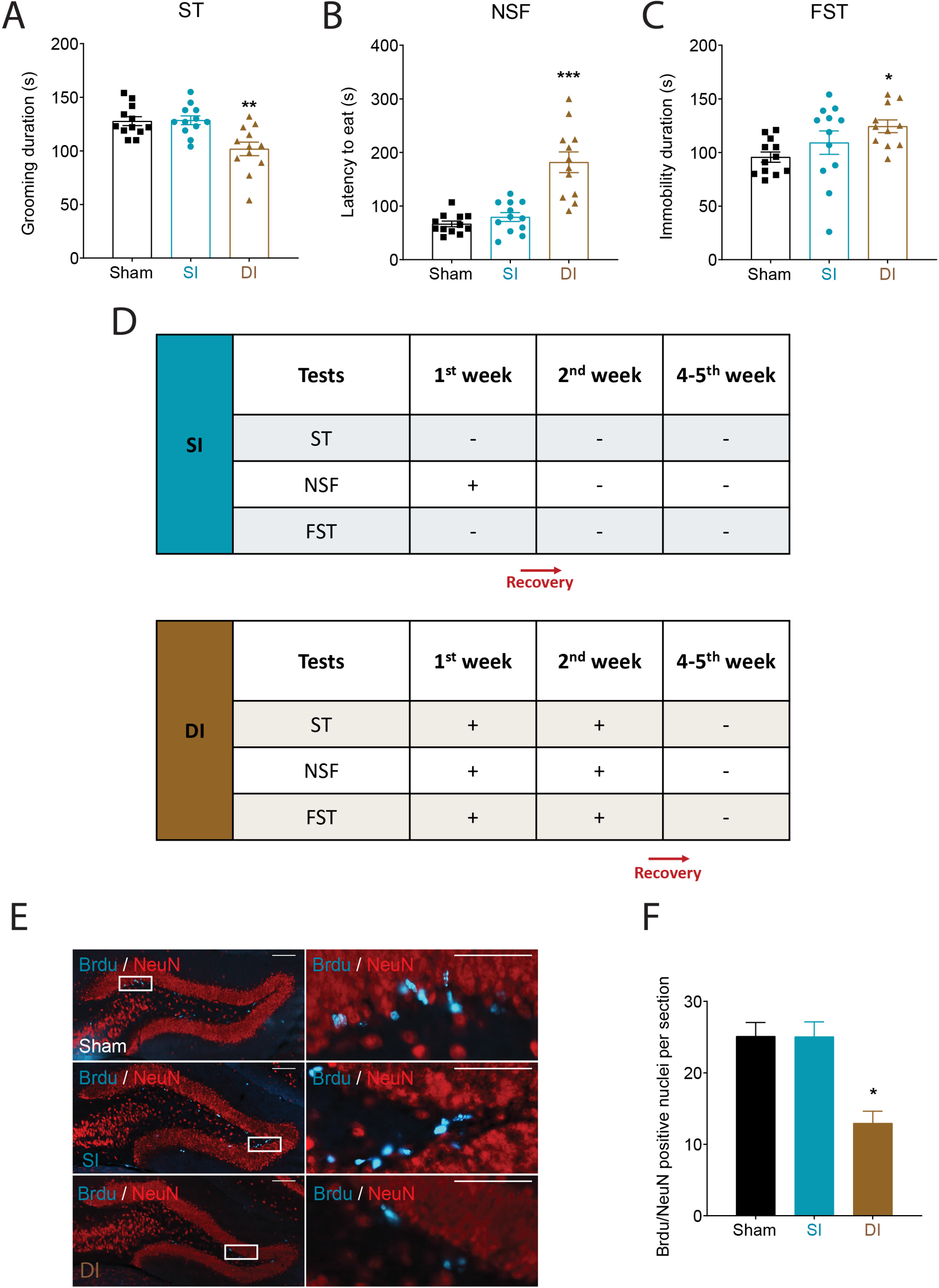
Influence of single (SI) or double incision (DI) on emotional behaviors and neurogenesis. On the second week after surgery (Sham, SI, DI), (**A)** decreased grooming behavior in the splash test, (**B**) increased latency to eat in the novelty suppressed feeding test and (**C**) increased immobility duration in the forced swim test is observed only in DI mice compared to control mice. (**D**) Summary of anxiodepressive-like behaviors in all tests at three different time points. (**E**) Brdu/NeuN positive nuclei per section in the dentate gyrus is significantly reduced only in DI mice 7 days post-injury (scale bars = 200μm) and (**F**) related quantification. All the values are means ± s.e.m. (n=12 except in **E, F**, n=4). One-way ANOVA and Holm-Sidak’s test (**A, B, C**) or Kruskal-Wallis test (**F**); *P<0.05; **P<0.01; ***P<0.001 vs. Sham.

### Characterization of pain after repeated hindpaw injuries

We also tested the effect of SI vs. DI on sensory thresholds and ongoing pain. Six hours after the first surgery, mice mechanical thresholds as measured by the VF test strongly decreased up to ten-fold in ipsilateral hindpaw (left hindpaw) indicating the appearance of mechanical allodynia (Fig. 2A). Mice recovered a normal mechanical threshold after 7 days, and mechanical threshold was unchanged in the SI contralateral hindpaw (right hindpaw). By contrast to the first incision, the second incision induced a robust decrease of mice withdrawal thresholds at the ispilateral side (right hindpaw) and at the contralateral side (left hindpaw; Fig. 2A). As reported in a previous work [34], recovery to a normal threshold of both hindpaws was delayed by 7 days compared to the first incision, with a full recovery at day 22. Area Under Curve (AUC) that represents the period of mechanical hypersensitivity observed the days after incision on the ipsilateral paw to the first or the second incision further confirmed that DI mice exhibited exaggerated mechanical pain hypersensitivity compared to SI mice (Fig. 2B). Mice also displayed prolonged thermal pain hypersensitivity after the second incision compared to the first one, as evaluated with the HG test (Fig. 2C). To evaluate spontaneous pain we used CPP experiments 3 days after surgery (SI or DI) or sham [42]. Only the DI experimental group developed a place preference for the clonidine-paired chamber (Fig. 2D) indicative of significant ongoing pain after DI.

**Figure 2:**
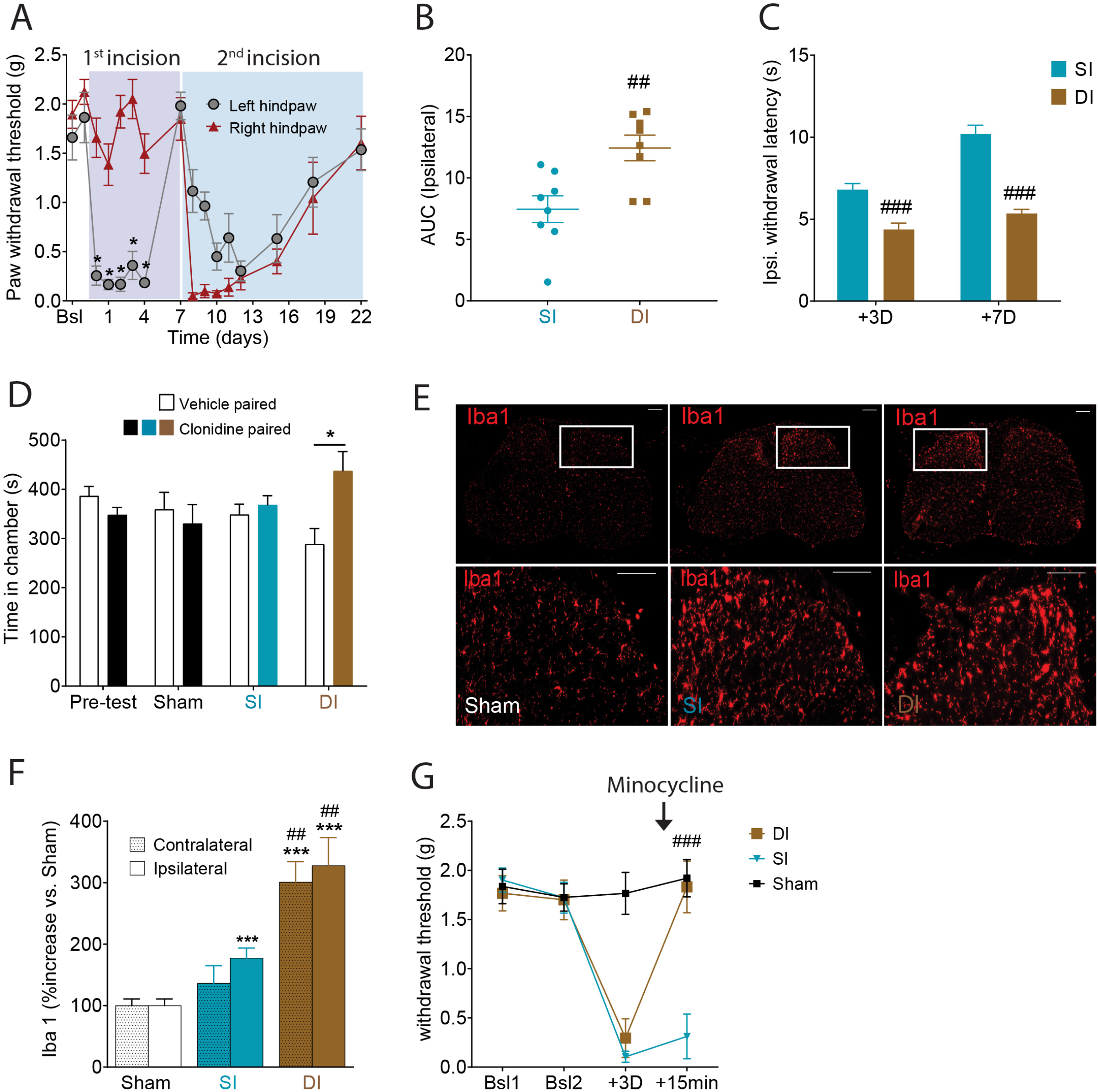
Influence of single (SI) or double (DI) incision on sensorial behaviors and microgliosis. (**A**) Mechanical hypersensitivity after incisions as measured by the Von Frey test and (**B**) cumulative area under curve representation show increased mechanical hypersensitivity after DI. (**C**) Ipsilateral withdrawal latency of Hargreave’s test shows increased thermal hypersensitivity after DI at D3 and D7. (**D**) Conditioned place preference induced by clonidine-evoked analgesia 4 days post-Sham, SI or DI showing a place preference only after DI. (**E**) Iba1 immunoreactivity in the whole spinal cord (left panel) and zoomed dorsal horn (right panel) 7 days after Sham, SI or DI (scale bars = 200μm) and (**F**) related quantification shows increased Iba1 staining density ipsilateraly after SI compared to sham, further increased after DI compared to SI. (**G**) Mechanical threshold of Sham, SI and DI animals before (3 days post-procedure) and 15 minutes after intrathecal injection of minocycline (300μg/mouse) show antinociceptive effect of minocycline only after DI. All the values are means ± s.e.m. (n=8 except in **D**, n=10 and **F-G**, n=4). Two-way ANOVA and Holm-Sidak’s test (**A**); Student’s t-test (**B-C**); One-way ANOVA and Holm-Sidak’s test (**D-G**); *P<0.05; **P<0.01; ***P<0.001 vs. Sham, baseline or vehicle paired; #P<0.05; ##P<0.01; ###P<0.001 vs. SI.

Next, to characterize the peripheral nervous system alterations induced by the procedures, we first topically applied Fluorogold to retrogradely label the DRG neurons affected by incision. At Day 7 (D7), most of the Fluorogold positive cells were found in L4 (DI: 27%; SI: 22%), few in L5 (DI, SI: ∼3%) and none in L6 DRGs (Fig. S2A-B). We then evaluated the expression of ATF3 and CSF1, two important molecular factors specifically expressed after neuronal stress or injury [43, 44],(Fig. S2 C-E). Similarly, at D7, most of ATF3 and CSF1 positive neurons were found in L4 (10-15%), few in L5 (∼3%) and 0% in L6 DRGs. We did not find any differential expression nor tracing differences between SI and DI mice suggesting that these peripheral alterations are unlikely responsible for the exaggerated pain-related behaviors after DI. Hence, we next examined, using real time quantitative PCR, the expression of different molecular and cellular markers involved in pain sensitization in the dorsal horn of the spinal cord. Interestingly, the nerve injury factors *Atf3* and *Sprr1a* mRNA in ipsilateral dorsal horn spinal cord were strongly increased after incision, with a potentiation in DI as compared to SI (Fig. S3A). We further quantified markers associated with microglial activation using RT-qPCR. Levels of Cd11b, Iba1, Csf1 and Csf1r mRNA were all found higher after incision with no differences between SI and DI conditions. We also looked at alterations of different spinal processes and cell-types through the labeling of peptidergic afferents (CGRP) and glial cells (Iba1 for microglia and GFAP for astrocytes). We found increased ipsilateral CGRP staining in dorsal horns of SI and DI compared to Sham mice at D3, without any difference between SI and DI mice (Fig. S3B-C). This increase was absent at D7 (Fig. S3D). While we found no difference in GFAP staining at D3 and D7 compared to the control group (Fig. S3E-G), Iba1 staining density was found increased compared to Sham at D7 (SI vs. Sham: x 1.8) and potentiated in DI compared to SI mice (DI vs. Sham: x 3.2; DI vs. SI: x 1.6) (Fig. 2E-F). To better understand the implication of microglia in the exaggeration of nociceptive behaviors, we then treated Sham, SI and DI mice with a single intrathecal injection of microglial inhibitors [45], minocycline (300μg/5μl/animal) at D3, when incision-induced pain hypersensitivity was still observed. Unexpectedly, minocycline restored mechanical thresholds to normal values only after DI and had no effect in Sham or SI mice (Fig. 2G). To test for a potential sex-related difference in microglia-mediated pain hypersensitivity as previously described in many reports [46–48], we repeated the experiment in both male and female mice after DI (Fig. S4). We observed a strong long lasting antinociceptive effect of minocycline in males still present 3 hours after injection. Yet, minocycline produced a weak and short-term antinociceptive effect in females (Fig. S4A). Note however that Iba1 staining density was still increased on the ipsilateral side (right hindpaw) in females after DI (Fig. S4B-C). To evaluate the effects of microglia proliferation in male animals, we used GW2580 (1μg/5μl/animal), which blocks microglia proliferation [49]. Similar results were obtained when intrathecally administered at the time of the surgeries (Fig. S5A-B) without affecting motor coordination (Fig. S5C). Strikingly, mice treated with GW2580 displayed enhanced grooming behaviors during the ST, reduced latency to eat during the NSF and immobility duration during the FST (P = 0.0570) compared to vehicle-treated mice indicating that the inhibition of spinal microgliosis also has a role in the development of mood disorders (Fig. S5D-F). Collectively we show here differentially expressed neuronal injury genes as well as increased microglia activation in the spinal cord after repeated injury bringing evidence of an increased central sensitization responsible for prolonged mechanical hypersensitivity and mood disorder comorbidity.

### Knocking-out FLT3 expression prevents both sensorial and emotional sensitization following DI

The overexpression of neuronal stress-related molecular markers (*Sprr1a* and *Atf3*) after DI also suggests a higher neuropathic component after DI compared to SI. We recently showed that peripheral FLT3 inhibition prevents the development of neuropathic pain [31]. Hence, we questioned the implication of FLT3 receptors in the development of exaggerated pain-related behaviors by assessing both sensorial and emotional dimensions after DI in *Flt3* Knock-out *(Flt3*^-/-^) female and male mice. In the VF test, the basal mechanical threshold was not affected by FLT3 silencing. After surgery, *Flt3*^-/-^ male mice displayed a faster recovery (Fig. 3A-C) along with reduced mechanical pain hypersensitivity to both the first and the second incision compared to male *Flt3* Wild Type (*Flt3*^+/+^) mice. Similar results were observed in female *Flt3*^-/-^ mice (Fig. S6A-B). CPP experiments also confirm the presence of a place preference for the clonidine-paired chamber in the DI *Flt3*^+/+^ group. This place preference was totally prevented in DI *Flt3*^-/-^ animals (Fig. 3D). To check for a potential role of FLT3 in basal mood modulation, we compared anxiodepressive-like behaviors of *Flt3*^-/-^ with *Flt3*^+/+^ animals in naive condition (Fig. S6C-E). We did not observe differences between *Flt3*^-/-^ and *Flt3*^+/+^ mice in the ST and NSF but *Flt3*^-/-^ already displayed decreased immobility duration in the FST. In addition, we evaluated the effects of FLT3 silencing on DI-induced anxiodepressive-like behaviors. Locomotor activity and motor coordination were assessed in all tested animals (Fig. 3E; Fig. S6F) and were found unchanged. *Flt3*^-/-^ animals showed no significant differences with WT mice in NSF but were significantly less immobile in the FST and presented increased grooming in the ST (Fig. 3G-I). To understand what could be the cellular alterations possibly responsible for the behavioral modifications after FLT3 inhibition, we thought to look at microglia activation. Iba1 staining in the dorsal horn of WT animals was increased while no change was found in *Flt3*^-/-^ animals after DI (Fig. 3I-L).

**Figure 3:**
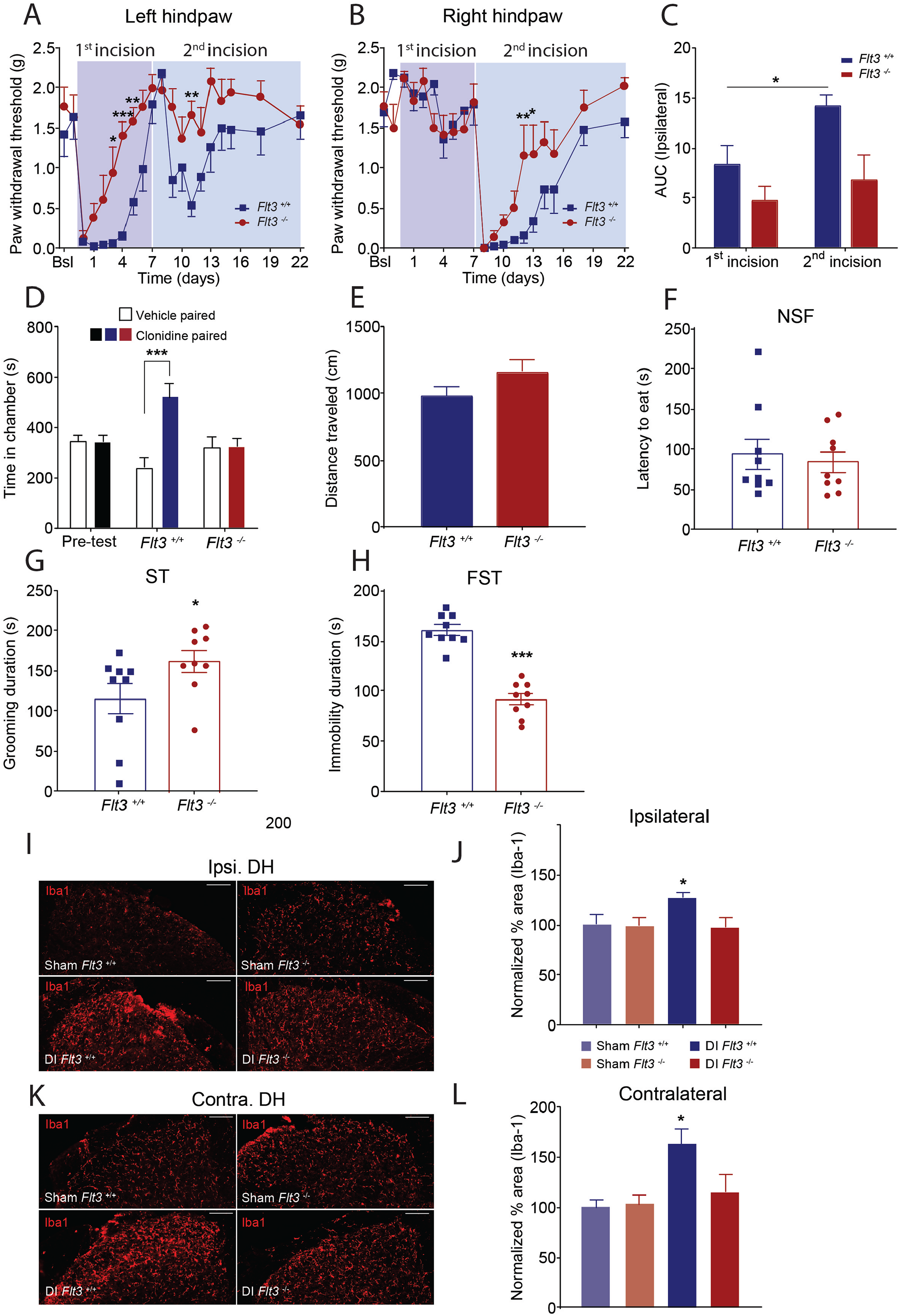
Silencing *Flt3* expression blocks DI-induced behavioral and molecular sensitization. (**A, B)** Left and right hindpaw mechanical hypersensitivity after incisions on *Flt3*^*+/+*^ or *Flt3*^*-/-*^ mice as measured by the Von Frey test and (**C**) cumulative area under curve representation highlight the absence of mechanical hypersensitivity exaggeration after DI in *Flt3*^*-/-*^. (**D)** Conditioned place preference induced by clonidine-evoked analgesia 4 days post-injury is totally prevented in *Flt3*^*-/-*^. Silencing *Flt3* expression does not change (**E**) locomotor activity and (**F**) latency to eat in novelty suppressed feeding test, but increases (**G**) grooming duration in splash test and decreases (**H**) immobility duration in forced swim test compared to *Flt3*^*+/+*^ mice. (**I)** Iba1 immunoreactivity of spinal cord ipsilateral dorsal horn dissected from Sham or DI (*Flt3*^*-/-*^ or control) mice and (**J**) related quantification show significantly higher labeling density in DI *Flt3*^*+/+*^ mice compared to Sham *Flt3*^*+/+*^. (**K**) Iba1 immunoreactivity of spinal cord contralateral dorsal horn dissected from Sham or DI (*Flt3*^*-/-*^ or control) mice and (**L**) related quantification show significantly higher labeling density in DI *Flt3*^*+/+*^ mice compared to Sham *Flt3*^*+/+*^. All the values are means ± s.e.m. (n=8/9 except in **I**-**L**, n=6/7). Two-way ANOVA and Holm-Sidak’s test (**A-B**); Student’s t-test (**E-H**); One-way ANOVA and Holm-Sidak’s test (**C, D, J, L**); *P<0.05; **P<0.01; ***P<0.001 vs. *Flt3*^*+/+*^ or Sham *Flt3*^*+/+*^ or Sham *Flt3*^*-/-*^.

### FL injection produces pain and mood disorders

To evaluate whether local FLT3 activation alone can induce both sensorial and emotional alterations as displayed in the DI model, mice received intrathecal injections of FL (50ng/5μl/animal) and were tested. As previously shown, a single injection led to decreased mechanical threshold for at least 2 days (Fig. 4A). Repeated intrathecal FL (2 weeks, 1inj/3days) failed to impair motor coordination but decreased grooming behavior, increased latency to eat and immobility duration on the ST, NSF and FST compared to vehicle-treated animals, respectively (Fig. 4B-E). All behaviors were significantly modified in all tests after repeated FL treatment, mimicking features of the DI model. At the cellular level, we previously reported that FL produced the activation in DRG of the stress-induced gene transcript Atf3 and several important neuronal pain-related gene transcripts [31]. Here, we show that Iba1 staining was found quantitatively increased after repeated FL injections (Fig. 4F-G), supporting a role for FLT3 in central changes. To rule out a possible implication of FLT3 expression in microglia cells after surgery, we performed Flt3 *in situ* hybridization of spinal cord from the CX3CR1^EGFP^ mouse line, in which EGFP is expressed in CX3CR1+ cells, known as macrophages (including microglia). Our data revealed an absence of colocalization between *Flt3* mRNA and CX3CR1 (Fig. S7). These data suggest that neuronal FLT3-induced spinal microglial activation promotes central pain sensitization and is associated with emotional and sensorial alterations.

**Figure 4:**
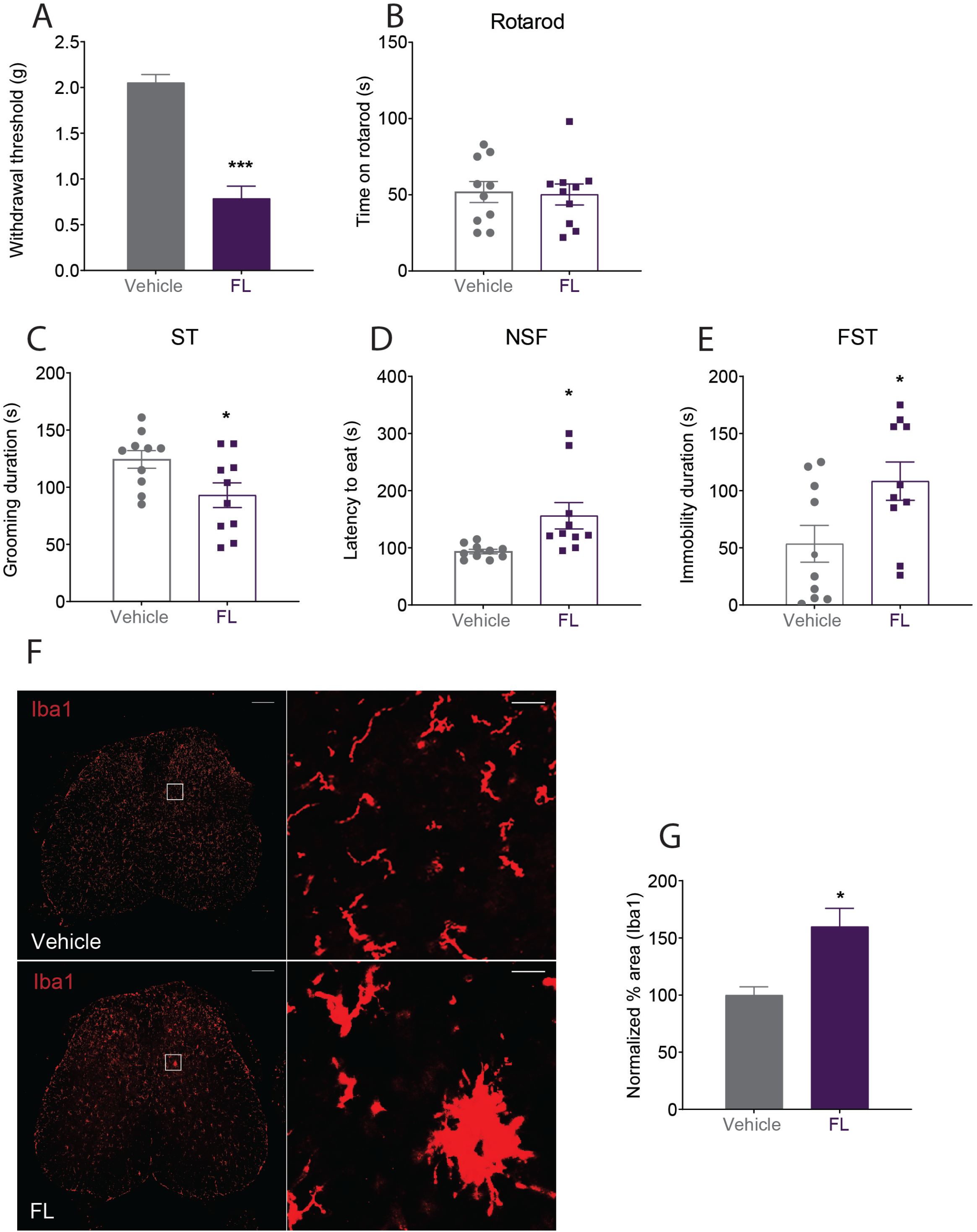
FL administration mimics DI induced behavioral and molecular alterations. (**A)** Mechanical hypersensitivity measured with Von Frey filaments after a single intrathecal injection of FL (50ng/5μl) or saline shows decreased withdrawal thresholds after FL. **(B)** Repeated intrathecal injections of FL (50ng/5μl, 5 injections delivered across 15 days) fail to affect performance on rotarod, (**C**) decrease grooming duration in splash test, (**D**) increase latency to eat in novelty suppressed feeding test and (**E**) immobility duration in forced swim test compared to control injections. (**F)** Iba1 immunoreactivity of whole spinal cord dissected from vehicle injected and FL injected mice (50ng/5μl, daily injection for 3 days) 24 hours after the last injection (left scale bars = 200μm; right scale bars = 20μm) and (**G**) related quantification shows increased Iba1 staining density after FL. All the values are means ± s.e.m. (n=10 except in **F, G**, n=4). Student’s t-test (**A-E**); unpaired Mann-Whitney t-test (**G**); *P<0.05; ***P<0.001 vs. I.t. Vehicle.

### Humanized antibodies against both human and mice FLT3 as a tool for conditional inhibition

Lastly, we developed cross-reacting antibodies specifically directed against both human and mice FLT3. Functional antibodies have the interesting property to not cross the blood brain barrier and could be an advantageous tool for questioning peripheral FLT3 function, and potentially treat postoperative pain in humans without CNS secondary effects. We produced the mAbA3 antibody that was found to present high affinity for both receptors (Fig. 5A-B; Fig. S8). Moreover, Homogeneous Time Resolved Fluorescence (HTRF) signals of FLT3-expressing RS4-11 cells in the absence or presence of FL were strongly reduced in a dose dependent manner after exposure to mAbA3 (Fig. 5C-E). Because we know the therapeutic benefits of inhibiting FLT3 in neuropathic pain models, we validated the *in vivo* efficacy of the antibody in the Chronic Constriction Injury (CCI) of the sciatic nerve model of neuropathic pain. Single or repeated (Fig. 6A, C) systemic injections of mAbA3 totally blocked mechanical pain hypersensitivity evaluated on the injured hindpaw. Inhibition of mechanical hypersensitivity started at a dose of 50 μg/animal of mAbA3 (Fig. 6B). We next tested the efficacy of mAbA3 in animals that underwent repeated incisions. A single preventive systemic injection of 200 μg of mAbA3 during the first incision strongly accelerated recovery to a basal mechanical threshold after both the first and the second incision on the ipsilateral hindpaw (Fig. S9A, B) without affecting either motor coordination (Fig S9C) or DI-induced anxiodepressive-like behaviors (Fig. S9D-F). We then determined whether repeated administration of mAbA3 was effective in reducing both sensory and anxiodepressive-related behaviors. Repeating preventive injections of mAbA3 (1 injection every 2 days starting from the day of the first incision until the second incision) not only reduced pain hypersensitivity but also totally prevented the development of anxiodepressive-like behaviors (Fig. 6D-H).

**Figure 5:**
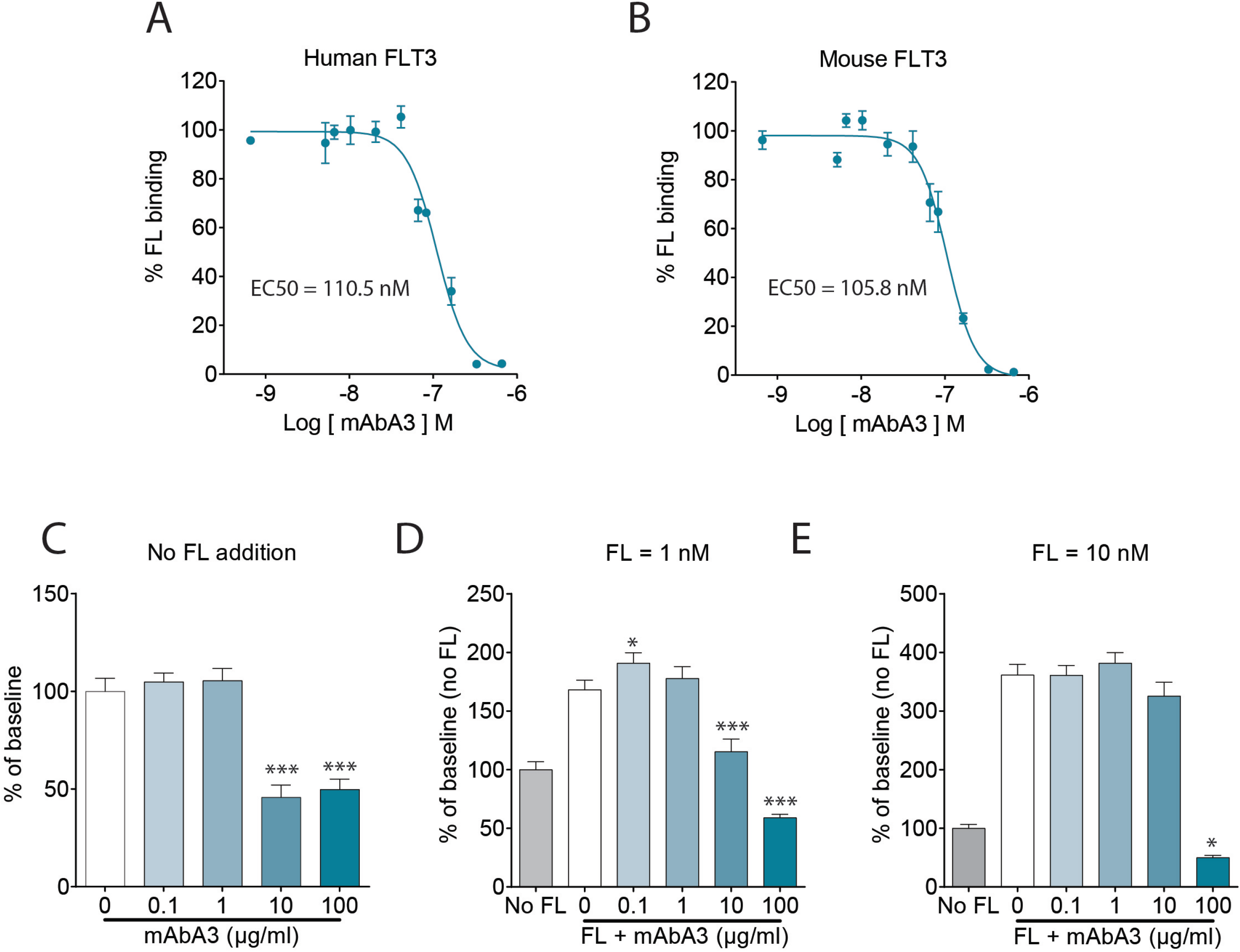
MAbA3 is a high affinity functional antibody against both human and murine FLT3 receptors. Binding curves of FL on human (**A**) and murine FLT3 (**B**) in presence of increasing doses of mAbA3. MAbA3 dose dependently reduces HTRF signal reporting mouse FLT3 constitutive activity (**C**) and FL-induced increased FLT3 activity after FL application at 1 nM (**D**) and 10 nM (**E)**. All the values are means ± s.e.m. (n=4); *P<0.05; **P<0.01; ***P<0.001 vs. Baseline

**Figure 6:**
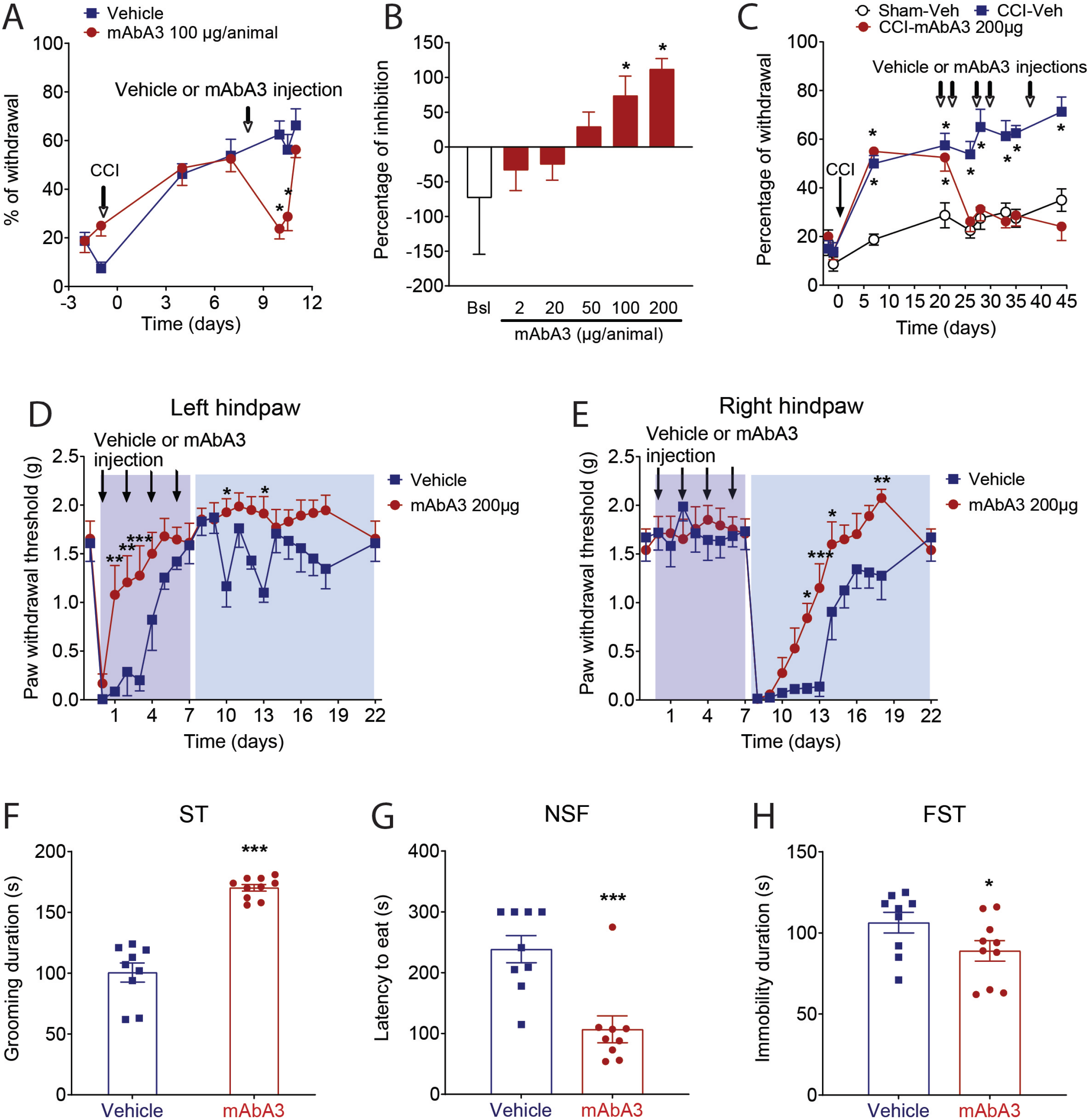
MAbA3 treatment against FLT3 prevents the development of CPSP. **(A)** A single injection of mAbA3 (100μg/mouse, intraperitoneal) reduces CCI-related (9 days after chronic constriction injury of the sciatic nerve) mechanical hypersensitivity as measured 1, 1.5 and 2 days after injection in the Von Frey test. (**B)** Experiments were performed as in **A**, with doses of mAbA3 ranging from 2 to 200μg/animal. Data represent the maximum percentage of pain hypersensitivity inhibition. (**C)** Intraperitoneal vehicle or mAbA3 injections (200μg/animal) were initiated 25 days after CCI of the sciatic nerve and repeated at times indicated by arrows. Mechanical hypersensitivity was recorded 26, 28, 33, 35 and 44 days after CCI. (**D-E)** Left and right hindpaw mechanical hypersensitivity after incisions in either vehicle or mAbA3 treated (arrows= 200μg/animal, intraperitoneal) mice as measured by the Von Frey test. Repeated intraperitoneal injections of mAbA3 (200μg/animal, 1 injection each 2 days during the first incision phase for a total of 4 injections) increases grooming duration in splash test (**F**), decreases latency to eat in novelty suppressed feeding test (**G**) and immobility duration in forced swim test (**H**) in treated mice compared to control mice. Two-way ANOVA and Holm Sidak’s test (**A-E**), Student’s t-test (**F-H**); *P<0.05; **P<0.01; ***P<0.001 vs. Sham-Veh or Vehicle.

## Discussion

Our results show that the repetition of surgical incision (DI) leads to sustained depressive-like behaviors along with the exaggeration of nociceptive behaviors and the appearance of spontaneous pain recapitulating the different features of chronic pain in humans [50]. The altered behaviors are associated with the increased expression of neuropathic-related molecular markers, a potentiation of microglia activation in the spinal cord as well as decreased neurogenesis in the hippocampus supporting central alterations after DI. We also report the implication of FLT3 in sustained pain sensitization produced by repeated injury. Strikingly, FLT3 activation alone is sufficient to trigger microglial activation, pain and anxiodepressive-like behaviors observed after DI. Conversely, FLT3 deletion leads to the complete prevention of DI-related behavioral and molecular modifications. Finally, we developed an innovative therapeutic tool, the FLT3 human antibody (mAbA3), targeting both human and murine FLT3. Treatment with FLT3 antibodies not only blocked exaggerated pain behaviors produced by repeated surgical procedure but also the subsequent depressive-like behaviors.

To better examine the mechanisms involved in pain chronification after acute injury, we used an experimental procedure that consists of repeating a surgical incision on the opposite hindpaw of the same animals 7 days apart [34]. Here, our main objective was to determine whether repeated acute injury may produce pain sensitization peripherally and centrally. To address this question, we characterized the differences observed between SI and DI at the behavioral and molecular levels. As previously reported, the pain hypersensitivity produced by the second surgery is largely enhanced compared to the one produced by the first surgery as it takes 7 days for the pain hypersensitivity to resolve after SI, compared to 14 days after DI. Our data suggest that surgery, like inflammation, may produce a sustained sensitized state leading to increased vulnerability to develop persistent pain when sensitized individuals are challenged with subsequent nociceptive stimuli leading to exaggerated nociceptive behaviors[16, 51]. However, to our knowledge no study has evaluated the impact of such a repetition on the development of emotional disorders. Our data revealed that DI not only produces exaggerated pain behaviors but also leads to sustained anxiodepressive-like behaviors starting after the second injury and lasting 3 weeks post-injury. Of note, behavioral alterations are associated with reduced hippocampal neurogenesis which is consistent with the depressive phenotype described in both rodents and humans [52–55]. Interestingly, depressive-like behaviors were already reported in other pre-clinical models of pain but usually develop after 5-7 weeks following nerve injury [56–58]. These models allow to identify the neuronal circuits supporting comorbidity of persistent pain and mood disorders [59, 60]. The DI model could thus become an outstanding model to study chronic pain but also pain-induced depression due to the rapidity of the development of pain and depression comorbidities in this model.

As already proposed, the mechanical pain hypersensitivity exaggeration produced by the second surgery on the previously non-operated hindpaw (right hindpaw) strongly suggests that pain sensitization is mainly affecting the central nervous system [51]. The central origin is also supported by the pain hypersensitivity triggered in the contralateral hindpaw (left hindpaw) to the second surgery. This is further underpinned by the fact that we did not observe any difference in the expression of molecular markers (ATF3 and CSF1) between DI and SI in DRG. Furthermore, retrograde labeling experiments highlight the small proportion of neurons affected by incision, mainly located in L4 DRGs. The behavioral modifications after DI could result in the summation of peripheral nociceptive inputs from both sides reaching the spinal cord leading to increased central changes. We thus evaluated potential modifications in the spinal cord of these animals. We showed that microglia activation is significantly enhanced after DI compared to SI or control. Note however that this was only observed at the protein level and that transcriptional changes related to microglia activation were found unchanged. An explanation could be that once microglia is activated, transcriptional changes are no longer required to potentiate microglia reactivity. Instead, post-transcriptional modifications would occur and regulate microglia. This aspect remains to be clarified. In males, the curative inhibition of microglial activation via minocycline or GW2580 leads to a quick recovery of pain sensitivity only after DI. Our observation agrees with previous data reporting that primed spinal microglia by neonatal incision is involved in the persistent pain in re-incised adult rats [61]. Interestingly, the microglial inhibition via G2580 also reduces anxiodepressive-related behaviors, indicating that spinal microglial overactivation produced by repeated incisions may be responsible for both the sensorial and emotional alterations caused by DI. In females, microglia is also activated after DI but, in agreement with previous reports [46–48], microglia inhibition has a less pronounced curative effect on mechanical hypersensitivity. Concerning astroglial activation, GFAP expression was not different in the spinal cord after SI or DI, as opposed to what has been reported in rats [62]. We can arguably propose that astrogliosis potentially appears later than the investigated timeframe. Altogether, our results show the involvement of central pain sensitization mechanisms in the exaggeration of nociceptive behaviors, in accordance with other chronic pain models [60].

In neuropathic pain models, central sensitization is in part mediated by CSF1 release from sensory neurons of the DRG [43] that then reaches the spinal cord to activate microglial CSF1R. Several observations across the study are supporting a major neuropathic component in the hindpaw incision model, especially after DI. First, we discovered a peripheral activation of CSF1 after both SI and DI, which is absent in an inflammatory pain model [63]. Secondly, this activation was associated with an activation of ATF3 expression in the majority of CSF1+ sensory neurons, although more expressed than CSF1 itself. Likewise, we also observed the spinal expression of *Atf3* and *Sprr1a* mRNAs, two markers of nerve injury [44, 64]. More importantly, FLT3 inhibition has been shown to induce therapeutic effects specifically in neuropathic pain rather than inflammatory pain models. The observation of a better post-surgical recovery after FLT3 inhibition brings more evidence of a neuropathic component inherent to our models. Finally, the increased expression of the spinal markers in DI vs. SI suggests a stronger neuropathic component in DI compared to SI supporting the clinical observation that persistent post-surgical pain is partially neuropathic in nature [65].

In the continuity of our previous work showing the involvement of FLT3 in neuropathic pain and considering the important neuropathic contribution in these models, we then evaluated the effects of FLT3 inhibition and activation in incision-induced central sensitization. Absence of *Flt3* expression in *Flt3*^-/-^ mutant mice leads to faster recovery of pain hypersensitivity after incision, and to a total prevention of spontaneous pain and anxiodepressive-like behaviors. Notably, beneficial behavioral effects of FLT3 silencing are common to both males and females. Spinal microgliosis activation is also reduced in *Flt3*^-/-^ animals. A major finding of the study is that beside the effects on DRG previously reported by our team [31], FL alone can also activate microglia in the dorsal horn of the spinal cord and recapitulate all the aspects of emotional and pain sensitization exhibited in DI. Interestingly, the effect of FL on FLT3 seems to be independent of microglia itself because *Flt3* mRNA is absent in CX3CR1 positive cells. Moreover, since *Flt3* mRNA is poorly expressed in the superficial dorsal horn layers of the spinal cord, it is tempting to speculate that the intrathecal injection of FL leads to the activation of peripheral FLT3, which then trigger the activation of spinal microglia through an indirect mechanism. Future studies will aim at better understanding the neuro-immune interactions responsible for this mechanism. To test for a possible role of peripheral FLT3 in central sensitization leading to emotional and sensorial comorbidities, we then developed functional antibodies (mAbA3) to reach high affinity and specificity for both mouse and human FLT3 using phage-display and a scFv synthetic library [66, 67]. Our data show that a single injection of mAbA3 is sufficient to prevent surgery-induced exaggerated pain hypersensitivity but is insufficient to prevent the development of surgery-induced anxiodepressive disorders. We thus considered the repeated administration of antibodies starting before the first surgery and continued every two days until the second surgical challenge. The preventive use of mAbA3 also largely decreases the time of recovery and totally hinders the development of anxiodepressive disorders. These results clearly support the role of peripheral neuronal FLT3 in the development of pain sensitization after repeated injury, the crossing of functional antibodies into the central nervous system being impeded by the blood-brain barrier.

In conclusion, we demonstrate that repeated peripheral incisions lead to sustained sensorial and emotional alterations through central pain sensitization. Increased expression of axonal regeneration genes and markers of neuropathy unveils the strong neuropathic component in our models, particularly in DI. Our study supports peripheral FLT3 as an important upstream neuronal modulator of central pain sensitization leading to exaggerated pain-related behaviors and anxiodepressive disorders. Targeting FLT3 via functional inhibitory antibodies efficiently alleviates sensitization, opening new avenues for the management and the prevention of pain chronification and related mood disorders.

## Author contribution

CR, JV designed the project; AT, DG, IM, MT, AJ, CS, LD, JPL, SM did the experiments; MC, MN, BR, PM generated mAbA3; AT, DG analyzed the data; JV, CR provided resources; XC, CR directed the work; AT, CR and JV wrote the manuscript.

## Acknowledgments and Disclosure

This work was supported by the University of Montpellier and grants from the READYNOV Région Occitanie and European community program, SFETD (Gisèle Guilbaud 2019 price), CBS2 doctoral school and BIODOL therapeutics. We are grateful to F. Perrin for providing CX3CR1^EGFP^ mouse line, to B. Pau to help for designing FLT3 antibody strategy, to P. Sokoloff, P. Carroll and I. Yalcin for constructive comments on the manuscript. MAbA3 was produced by GenAc platform funded by the French National Research Agency under the program “Investissements d’avenir” Grant Agreement LabEx MAbImprove: ANR-10-LABX-53 and by the SIRIC Montpellier-Cancer under Grant Agreement “INCa-DGOS-Inserm 6045.

## Conflict of interest statement

C.S., L.D., and J.P.L. were full-time employees at Biodol Therapeutics. J.V is inventor of patents claiming the use of FLT3 inhibitors for the treatment of neuropathic pain and is co-founder of Biodol Therapeutics. The other authors declare no conflict of interests.

Supplementary information is available at MP’s website.

---

**Figure S1.**
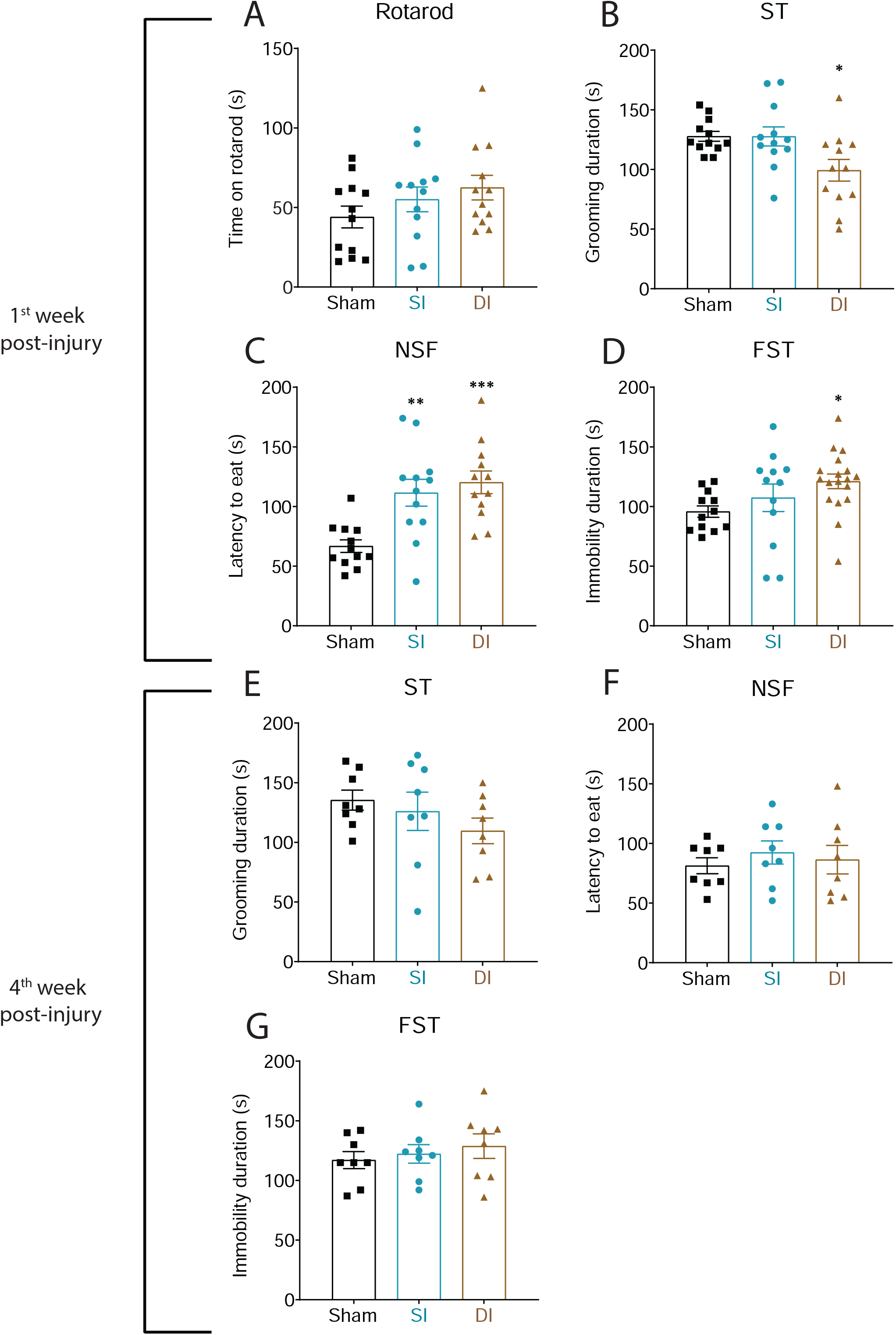

**Figure S2.**
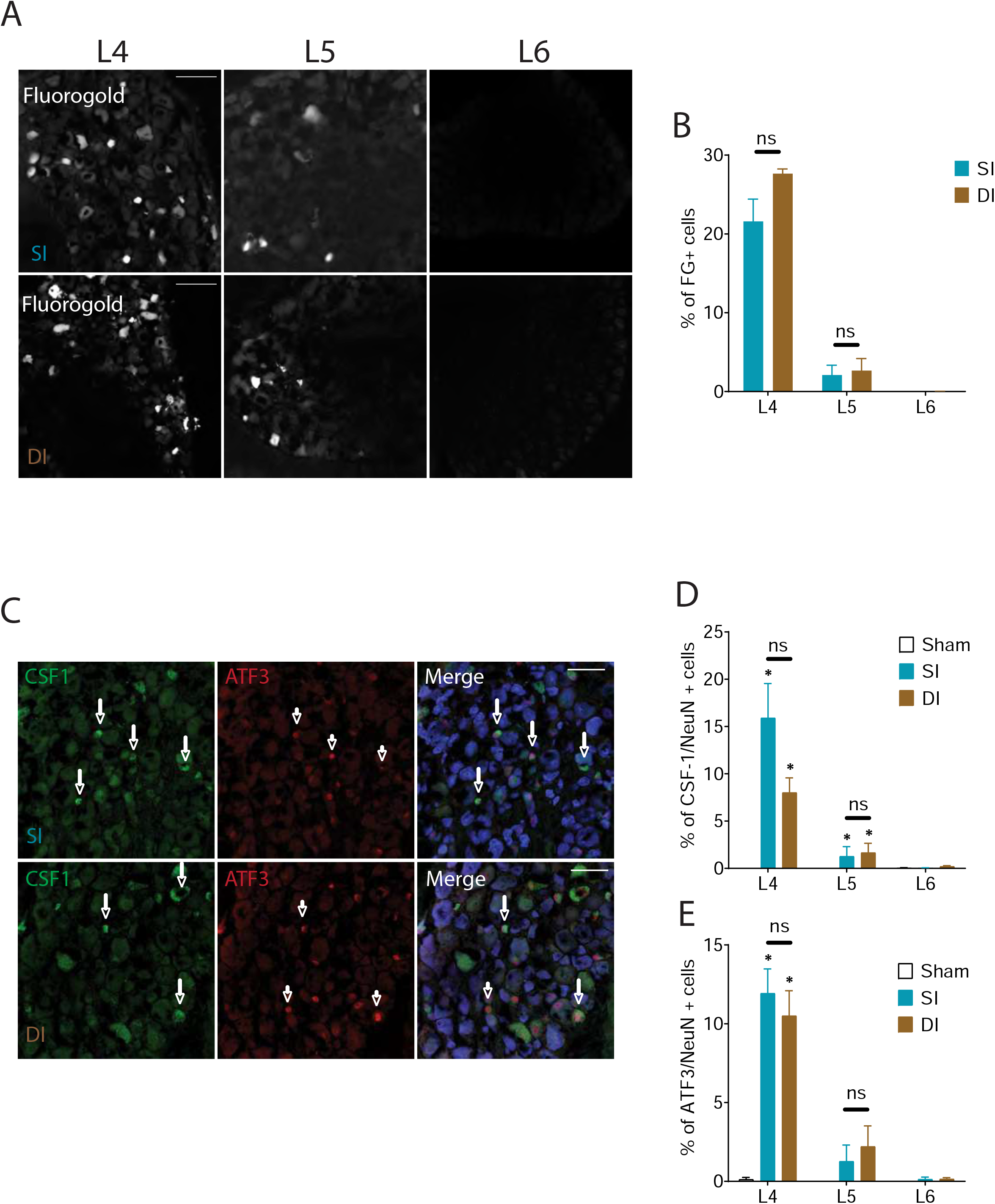

**Figure S3.**
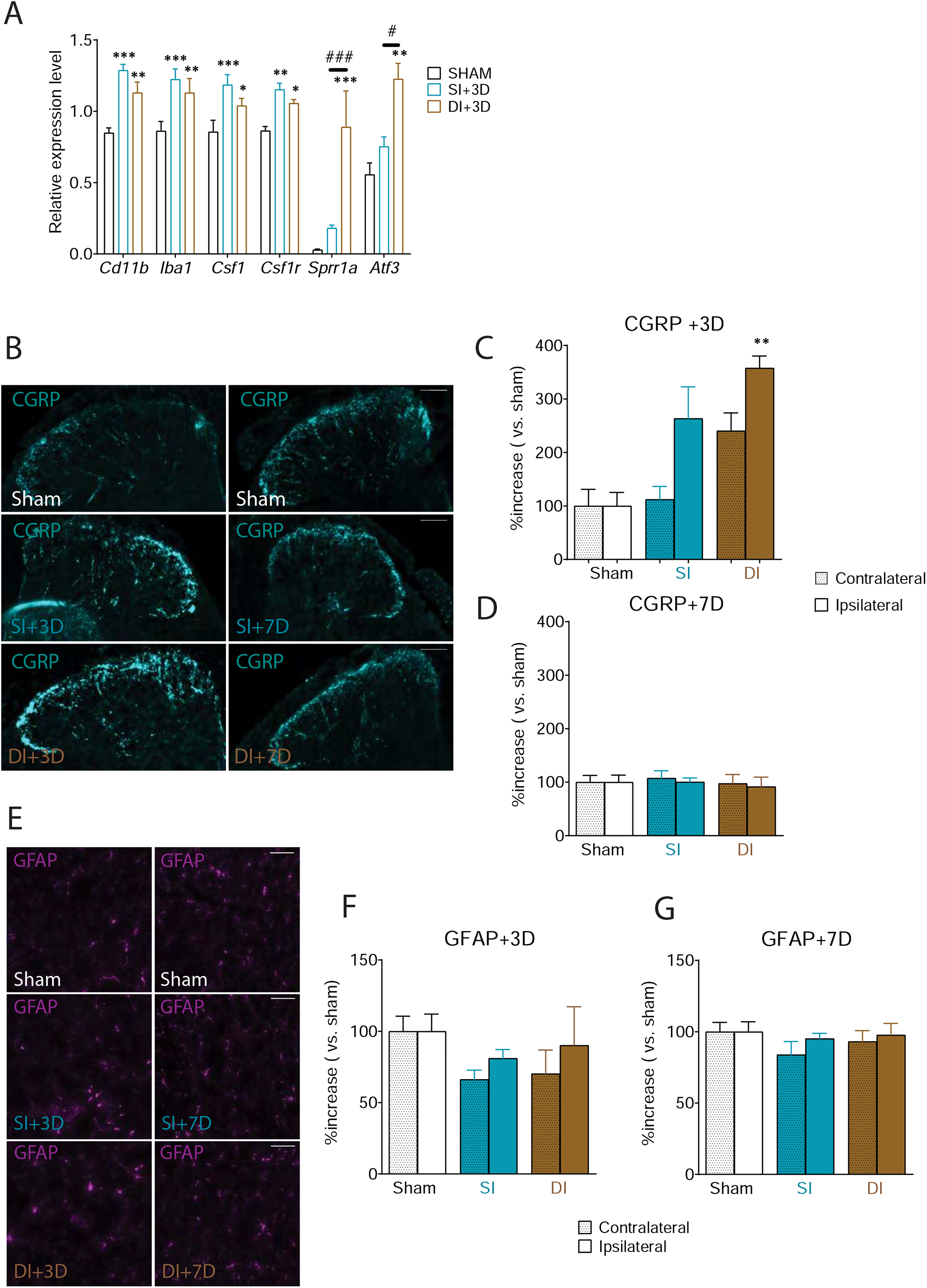

**Figure S4.**
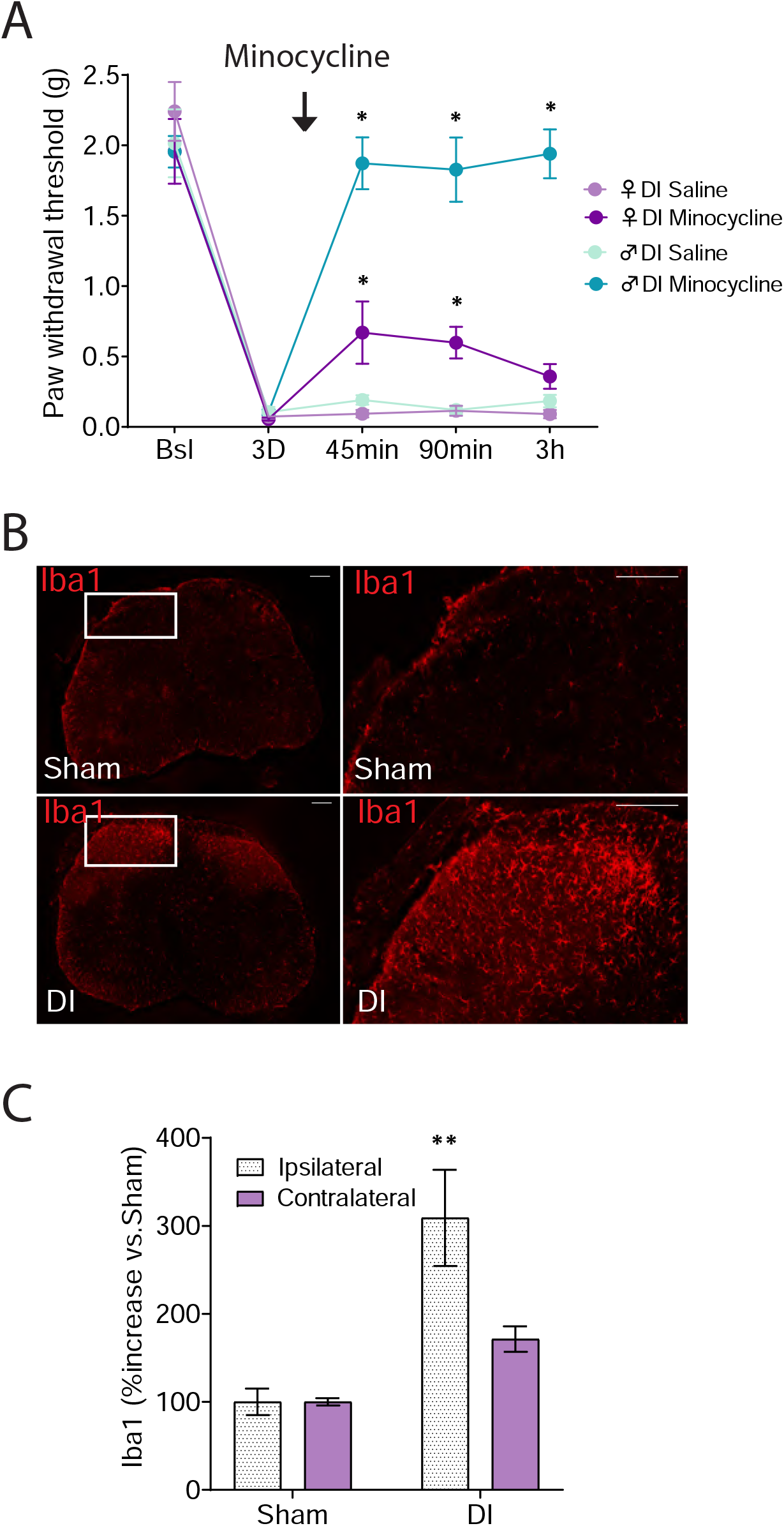

**Figure S5.**
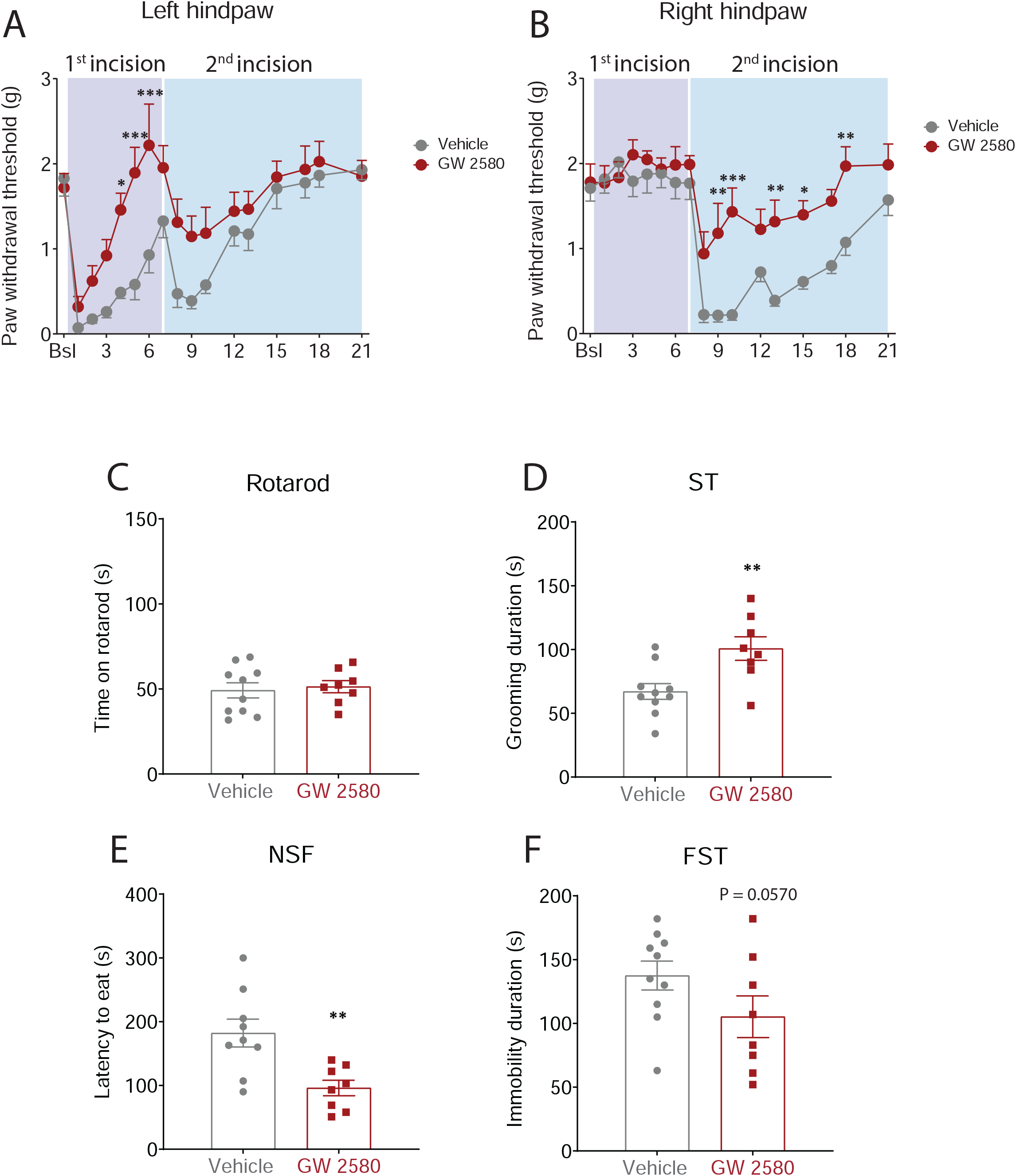

**Figure S6.**
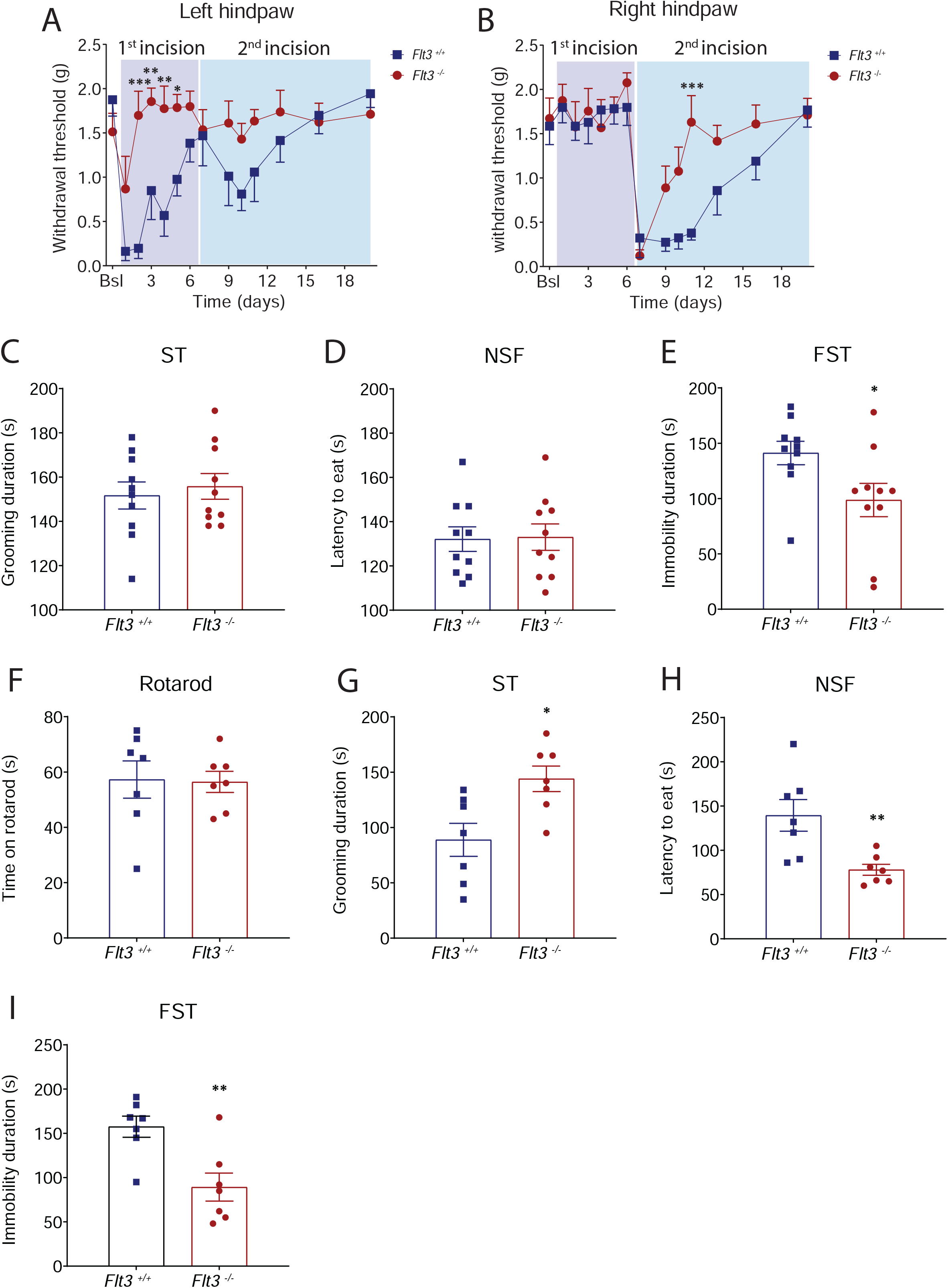

**Figure S7.**
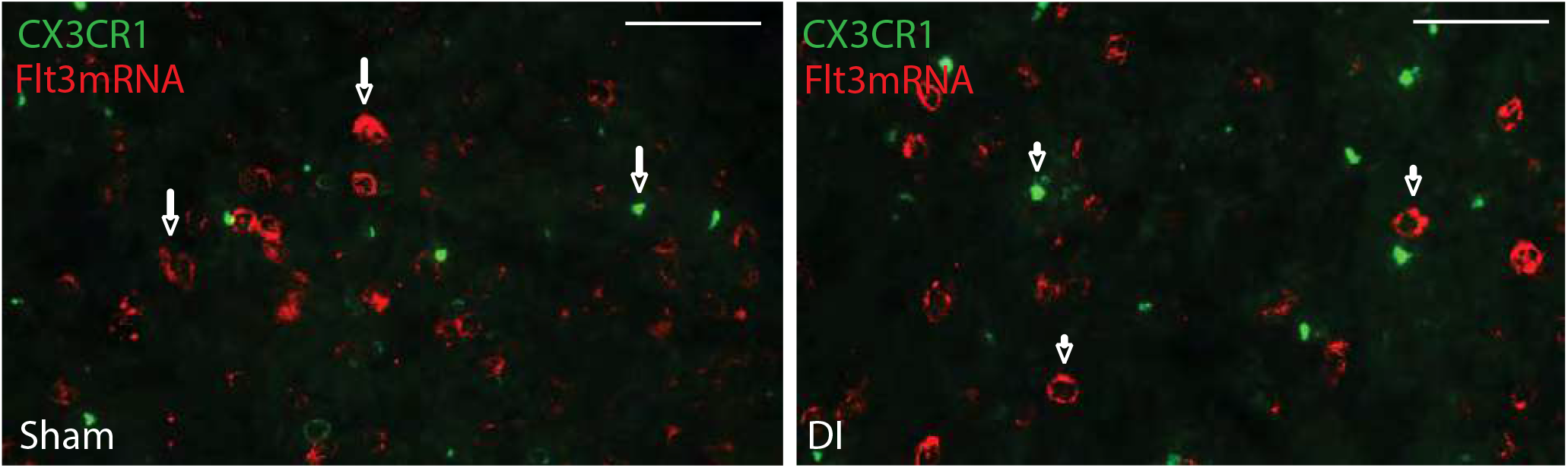

**Figure S8.**
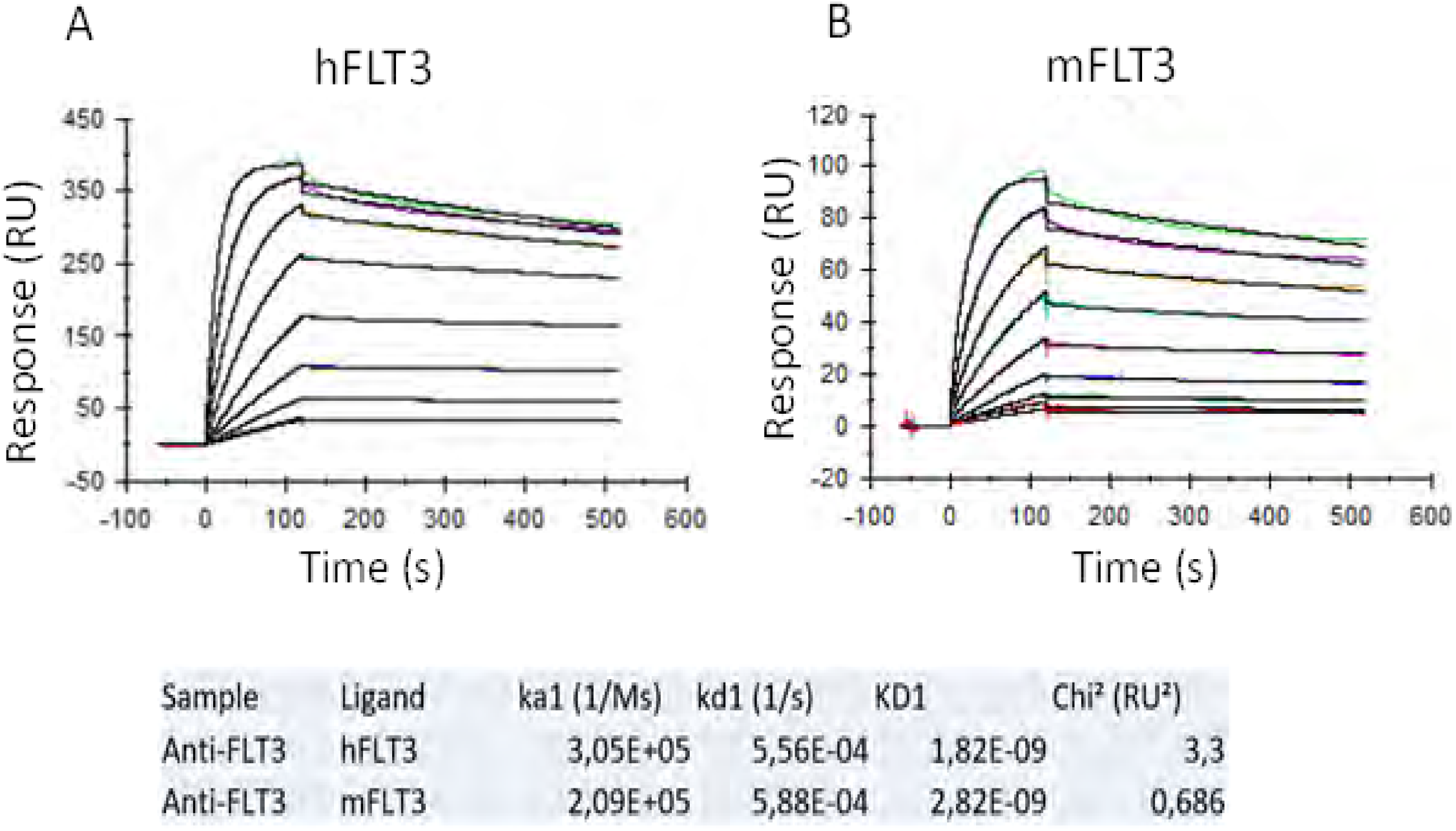

**Figure S9.**
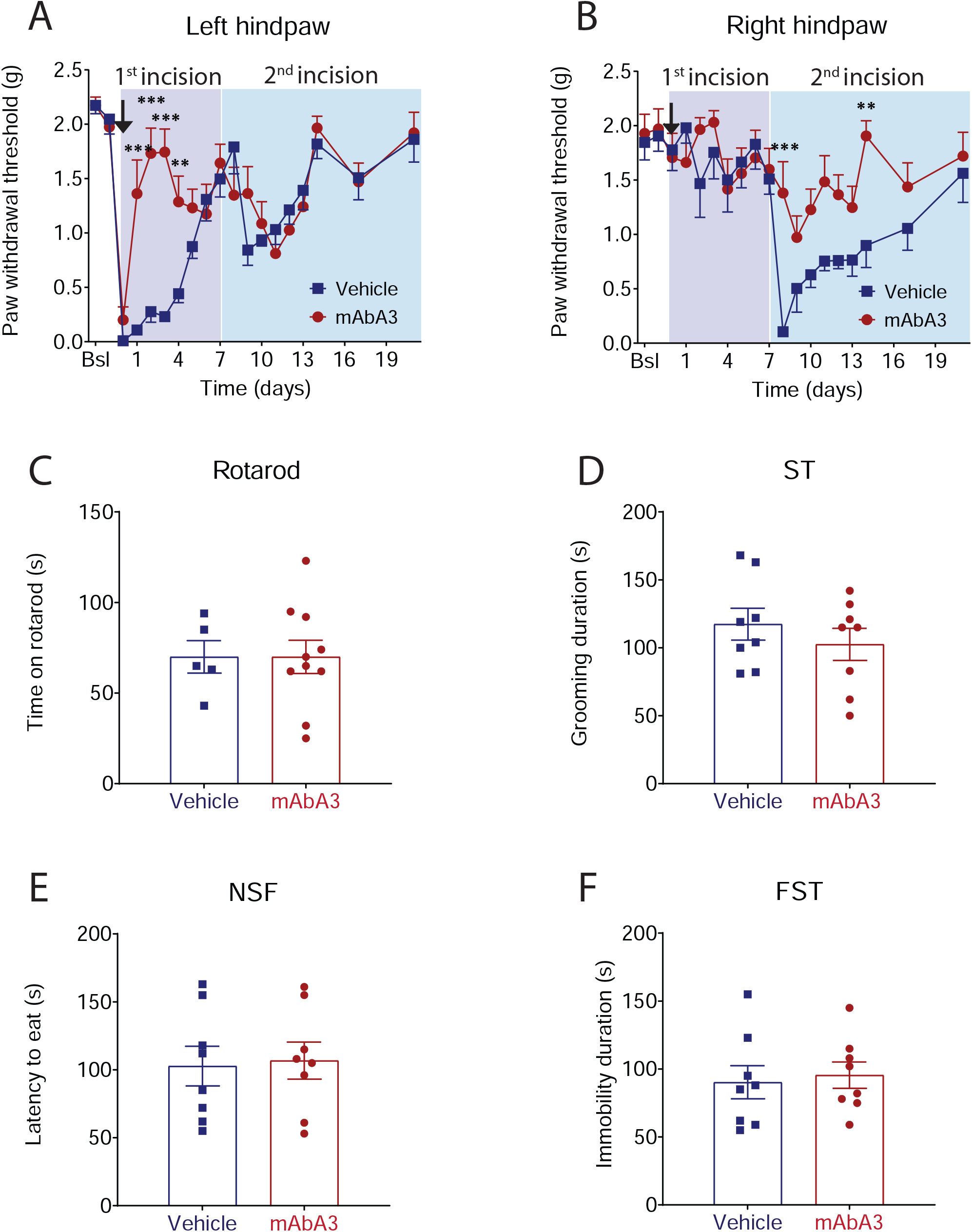

