## Supplementary file for "Activation of neuronal FLT3 promotes exaggerated sensorial and emotional pain-related behaviors facilitating the transition from acute to chronic pain"

Adresses : [1] Univ Montpellier, Montpellier, France ; [2] Inserm U-1298, Institut des Neurosciences de Montpellier, Montpellier, France ; [3] CNRS UMR 5203, Institut de Génomique Fonctionnelle, Montpellier, France ; [4] Département d'anesthésiologie, Hôpital Universitaire Lapeyronie, Montpellier, France ; [5] IRCM, INSERM U1194, ICM, Montpellier, F-34298, France, [6] BIODOL Therapeutics, Cap Alpha, Clapiers, France.

$ current address: University of North Carolina at Chapel Hill, Neuroscience center, Mary Ellen Jones, building, Chapel Hill, NC 27599, USA

**Supplementary figures: 9**

**Supplementary tables: 2**

**Supplementary methods and materials: 1**

### Supplementary figures

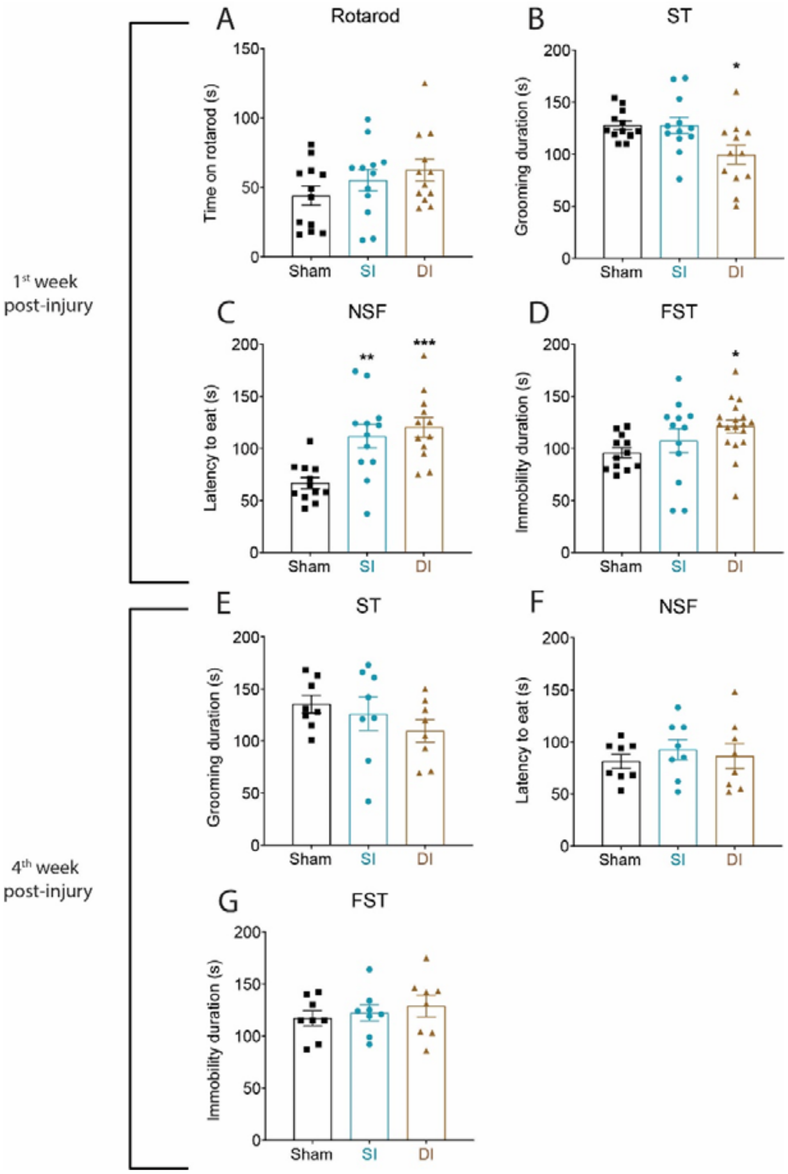


**Supplementary figure S1:** Influence of single (SI) or double incision (DI) on emotional behaviors. During the first week post-injury, (**A)** Neither SI nor DI significantly reduces performance in the rotarod test. (**B)** Decreased grooming behavior in the splash test (ST), (**C**) increased latency to eat in the novelty suppressed feeding test (NSF) and (**D**) increased immobility duration in the forced swim test (FST) is observed in DI and SI mice compared to control mice. In the fourth week, (**E**) no significant changes were found in the splash test, (**F**) novelty suppressed feeding test and (**G**) forced swim test in DI mice compared with control mice. All the values are means ± s.e.m. (n=12). One-way ANOVA and Holm-Sidak's test; *P<0.05; **P<0.01; ***P<0.001 vs. Sham.

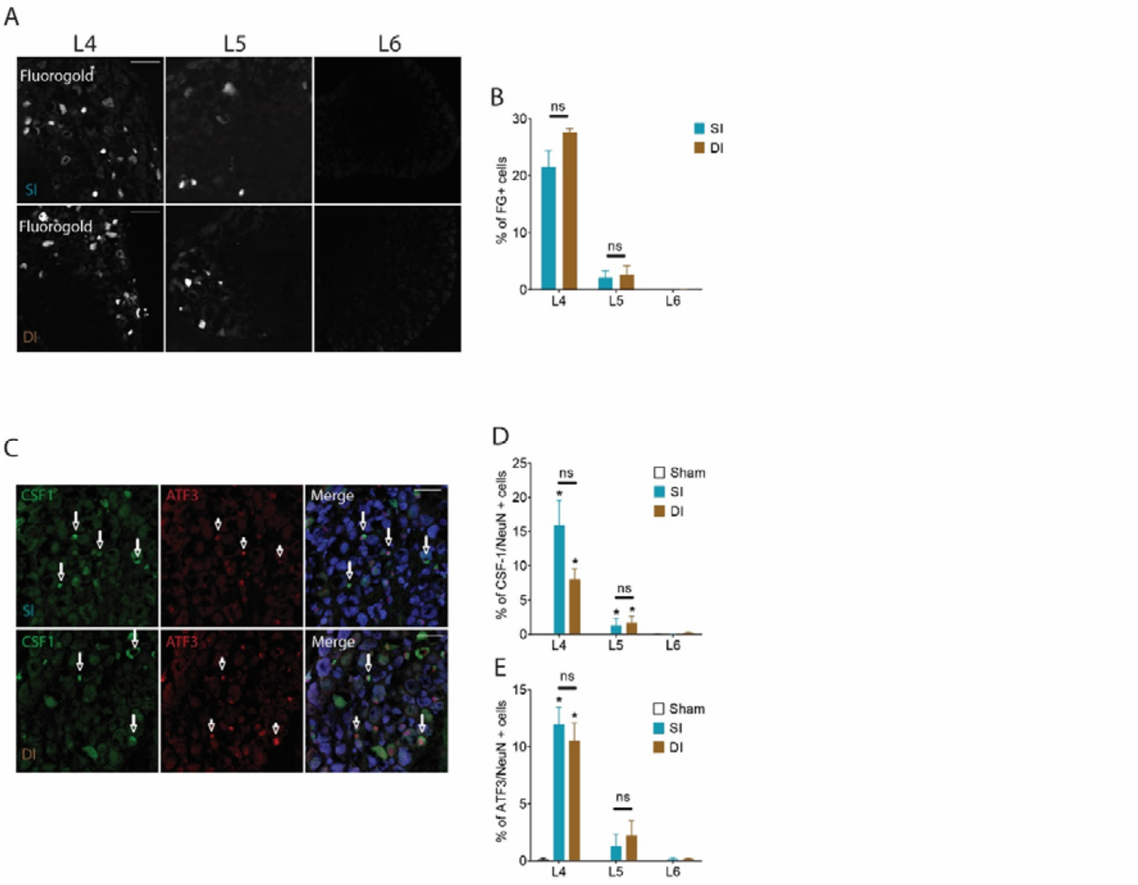


**Supplementary figure S2:** Influence of single (SI) or double incision (DI) on DRGs. (**A**) Fluorogold fluorescence in neurons of ipsilateral L4, L5 and L6 DRGs 7 days after SI or DI (scale bars = 100µm) and (**B**) related quantifications. **(C)** CSF1(long arrows) and ATF3 (short arrows) immunoreactivity in neurons of ipsilateral L4, L5 and L6 DRGs 7 days after SI and DI (merge = CSF1/ATF3/NeuN) and (**D, E**) related quantifications. All the values are means ± s.e.m. (n=8). One-way ANOVA and Holm-Sidak's test; ns: non-significant, *P<0.05 vs. Sham.

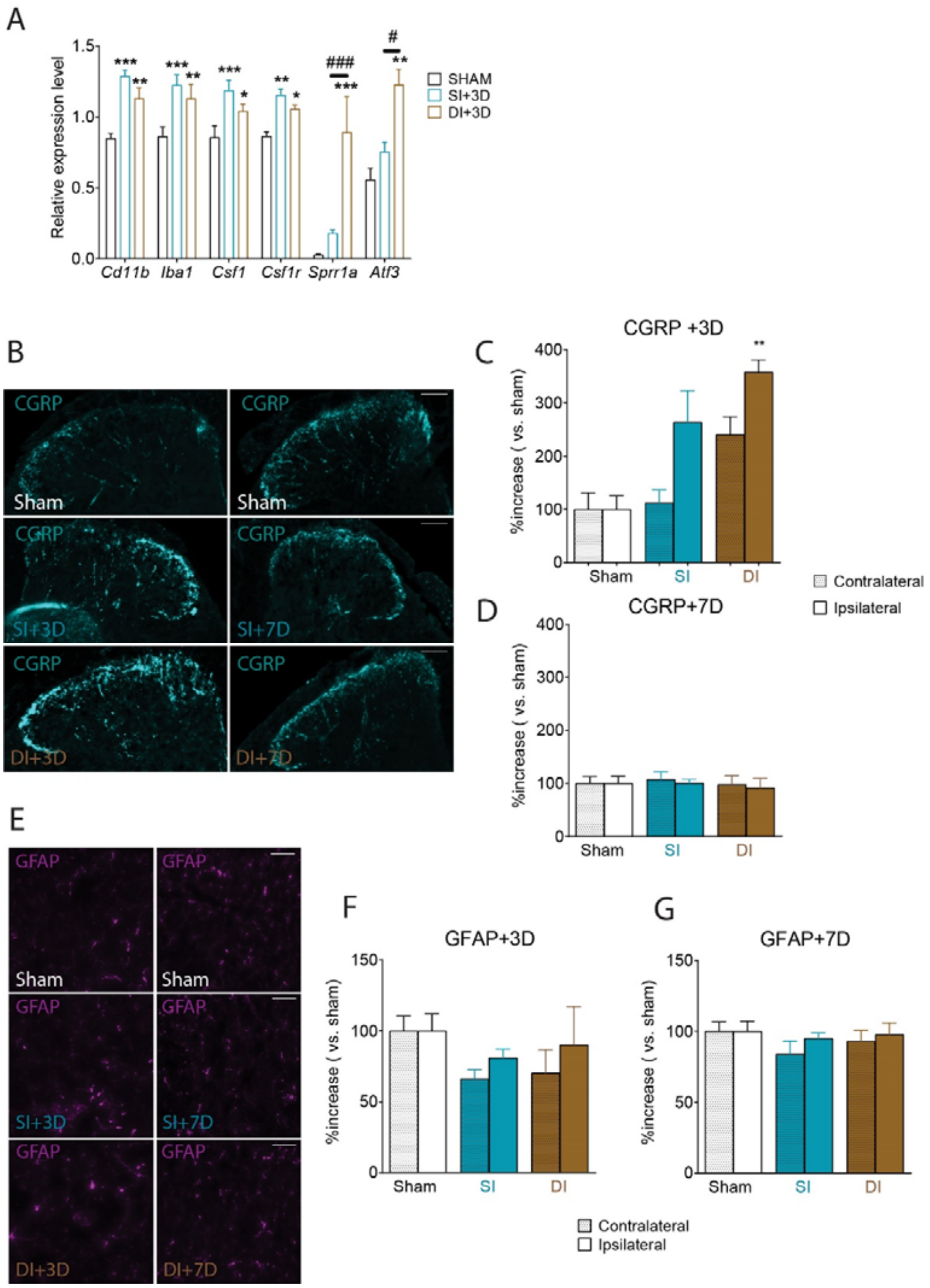


**Supplementary figure S3:** Influence of single (SI) or double incision (DI) on dorsal horn spinal cord cellular modifications. **(A)** Relative expression level of *Cd11b*, *Iba1*, *Csf1*, *Csf1r*, *Sprr1a* and *Atf3* mRNA in ipsilateral dorsal horn spinal cord 3 days after Sham, SI and DI. **(B)** CGRP immunoreactivity in dorsal horn spinal cord 3 and 7 days after Sham, SI and DI and (**C, D**) related quantifications. (**E**) GFAP immunoreactivity in dorsal horn spinal cord 3 and 7 days after Sham, SI and DI and (**F, G**) related quantifications. All the values are means ± s.e.m. (n=4). One-way ANOVA and Holm-Sidak's test; **P<0.05; ***P<0.05 vs Sham. #P<0.05; ###P<0.001 vs. SI.

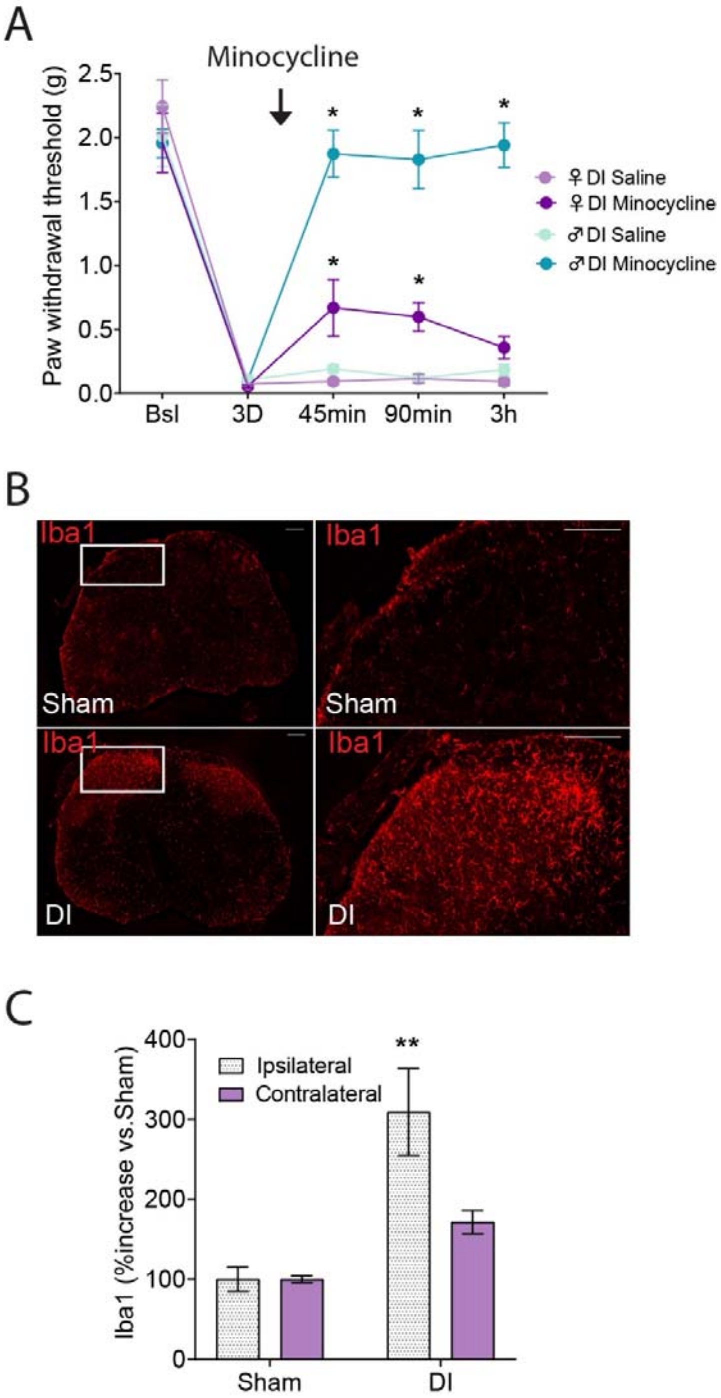


**Supplementary figure S4:** Microglia changes after DI. **(A)** Mechanical threshold of DI male and female animals before (3 days post-procedure) and 15 minutes after intrathecal injection of minocycline (300µg/mouse). **(B)** Iba1 immunoreactivity in the dorsal horn spinal cord 7 days after Sham or DI procedure in female mice (scale bars = 200µm) and **(C)** related quantification. All the values are means ± s.e.m. (n=4 except in **A**, n=8). One-way ANOVA and Holm-Sidak's test; **P<0.05; ***P<0.05 vs Sham.

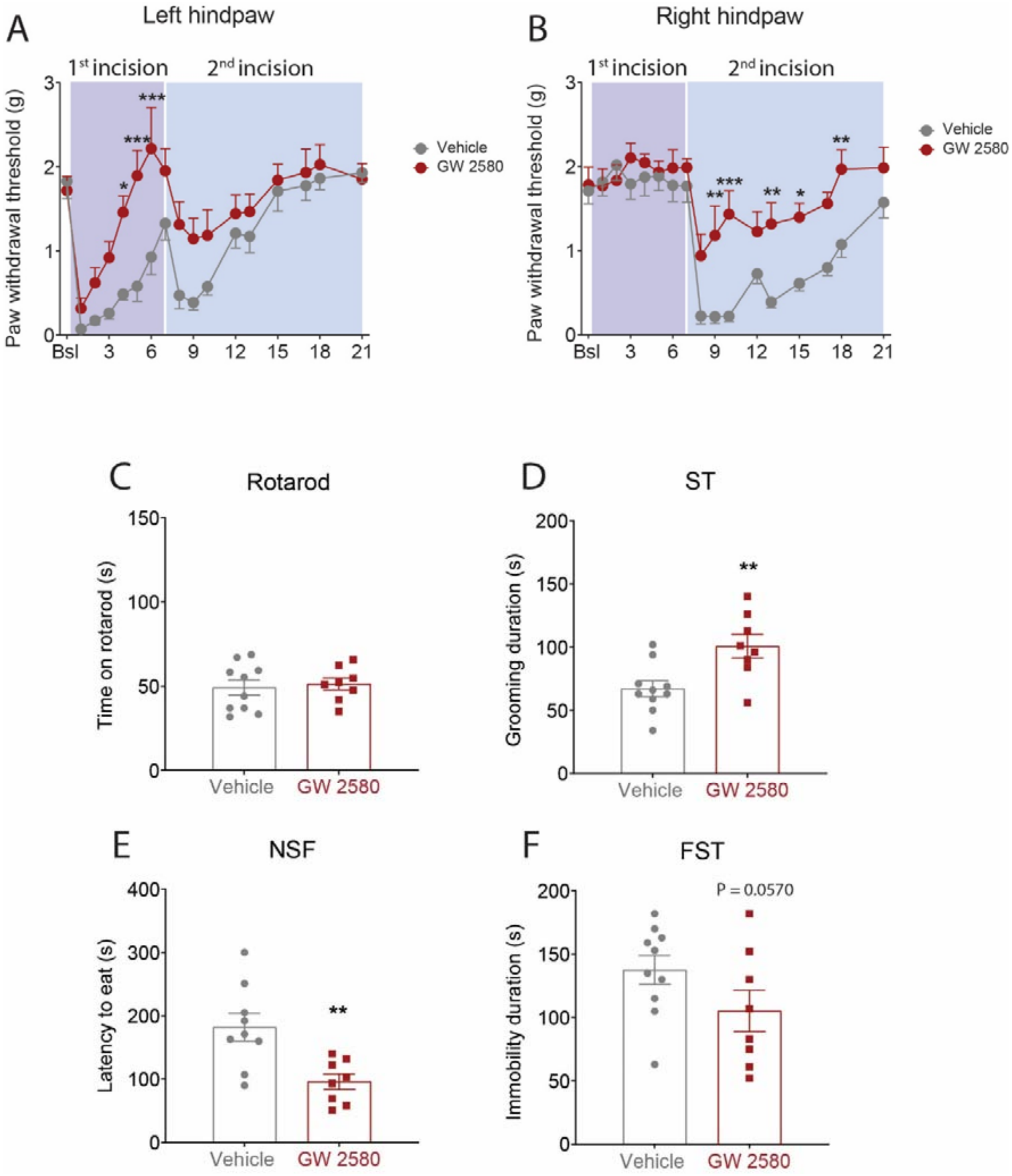


**Supplementary figure S5:** Effects of microgliosis inhibition on double incision (DI)-induced pain sensitization. **(A, B)** Intrathecal injections of GW 2580 (1µg/5µl, during each surgery) improve the recovery of a normal mechanical threshold after incision (SI or DI) as measured with Von Frey filaments. The same treatment fails to affect performance on rotarod (**C**), increases grooming duration in splash test (ST) (**D**), decreases latency to eat in novelty suppressed feeding test (NSF) (**E**) and decreases immobility duration in forced swim test (FST) compared to control injections (**F**). All the values are means ± s.e.m. (n=8). Two-way ANOVA and Holm Sidak's test (**A, B**), Student's t-test (**C-F**); *P<0.05; **P<0.01; ***P<0.001 vs. i.t. Vehicle.

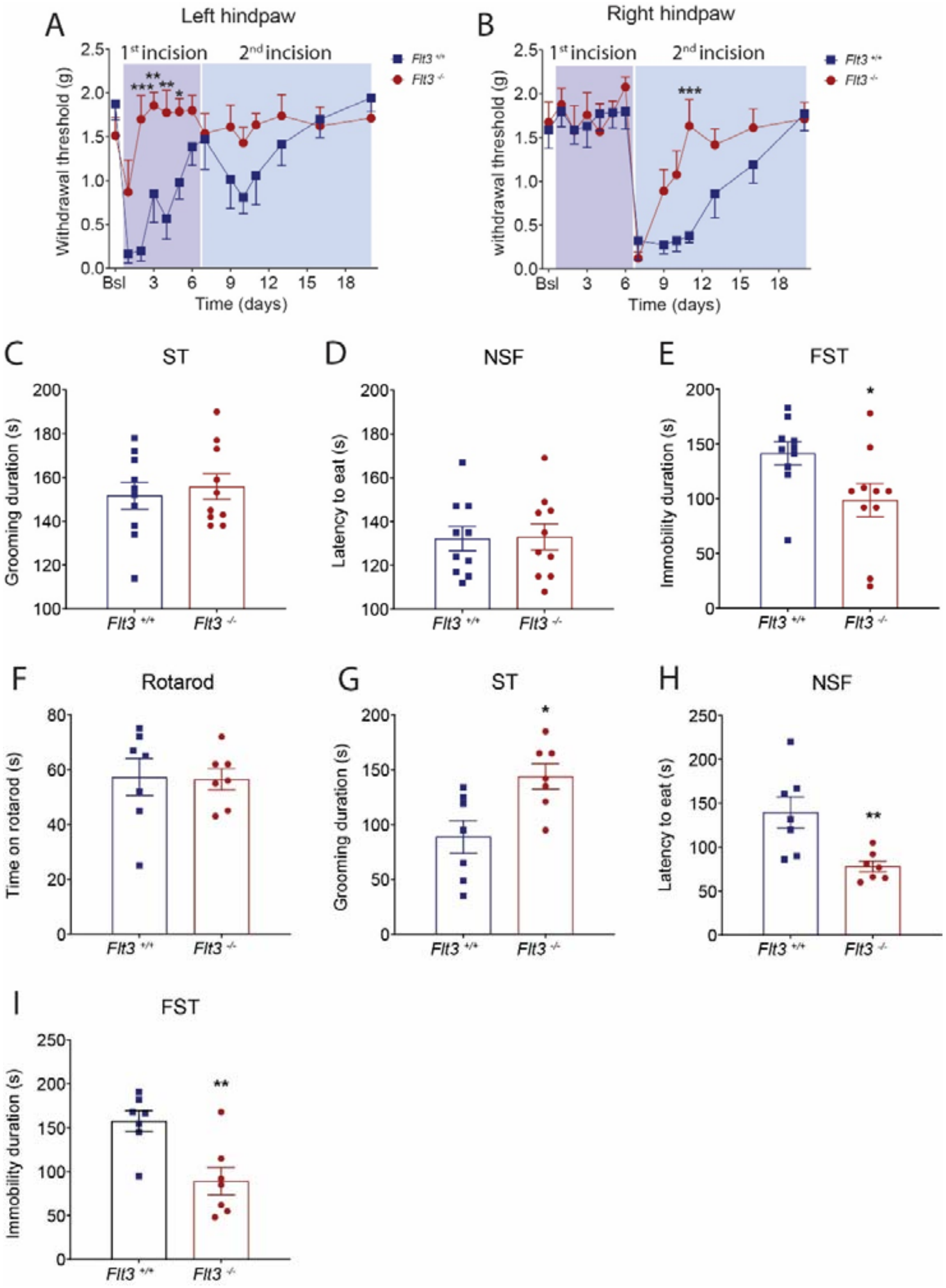


**Supplementary figure S6:** Behavioral phenotyping of *Flt3*$^{-/-}$ female mice before and after DI. (**A**) Genetic *Flt3* silencing does not affect grooming duration in the splash test (ST) and (**B**) latency to eat in the novelty suppressed feeding test (NSF), but (**C**) decreases immobility duration in the forced swim test (FST) compared to control mice. **(D-E)** Left and right hindpaw mechanical hypersensitivity after incisions in either *Flt3*$^{+/+}$ or *Flt3*$^{-/-}$ female mice as measured by the Von Frey test. **(F)** Genetic *Flt3* silencing in female mice fails to affect performance on rotarod, (**G**) increases grooming behavior, (**H**) decreases latency to eat in novelty suppressed feeding test and (**I**) immobility duration in forced swim test compared to control mice two weeks after DI. All the values are means ± s.e.m. (n=8). Two-way ANOVA and Holm Sidak's test (**D-E**), Student's t-test (**A-C; F-I**); *P<0.05; **P<0.01; ***P<0.001 vs. *Flt3*$^{+/+}$.

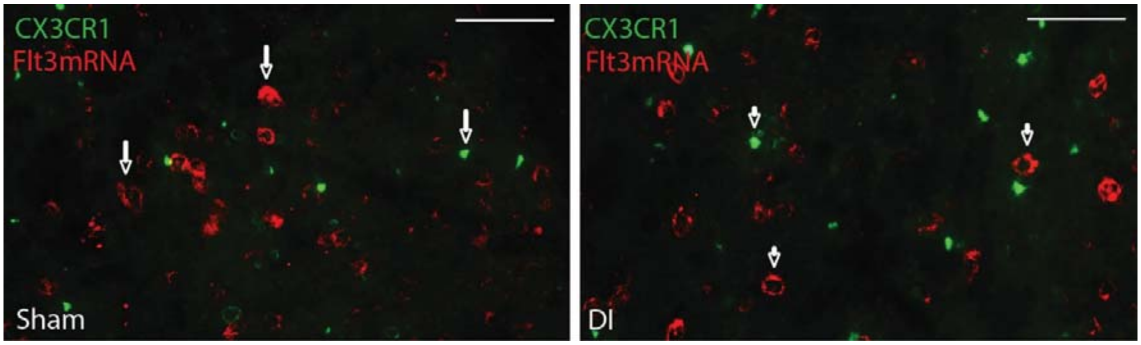


**Supplementary figure S7:** *Flt3* mRNA is not colocalized with CX3CR1. *In situ* hybridization of *Flt3* mRNA combined with immunoreactivity of CX3CR1 of spinal cord 7 days after Sham or DI. Cells from spinal cords are shown (scale bars = 100µm).

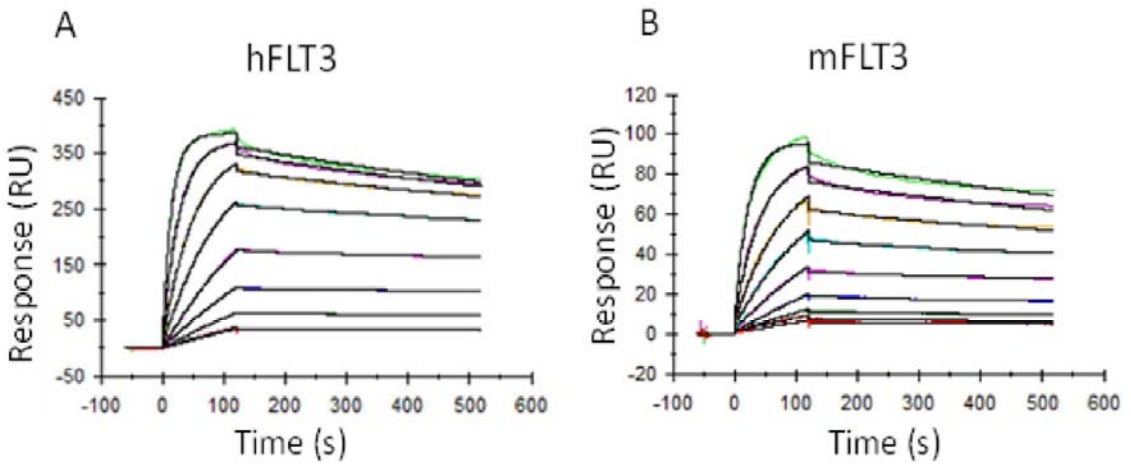


| Sample | Ligand | ka1 (1/Ms) | kd1 (1/s) | KD1 | Chi² (RU²) |
|---|---|---|---|---|---|
| Anti-FLT3 | hFLT3 | 3,05E+05 | 5,56E-04 | 1,82E-09 | 3,3 |
| Anti-FLT3 | mFLT3 | 2,09E+05 | 5,88E-04 | 2,82E-09 | 0,686 |

**Supplementary figure S8:** Affinity measurements of anti-FLT3 mAbA3 on human and murine FLT3- Fc-captured on CM5S sensor Chip using T200 BIAcore apparatus. Increasing concentrations (0.75nM, 1.5nM, 3.1 nM, 6.2nM, 12.5nM, 25nM, 50nM, 100 nM) of anti-FLT3 mAbA3 were injected on **(A)** human and **(B)** murine FLT3. Black curves are the fitting curves obtained by using a bivalent model. Association constant (ka1) and dissociation constant (kd1) of mAbA3 for human FLT3 are in a similar range than for murine FLT3 (Table).

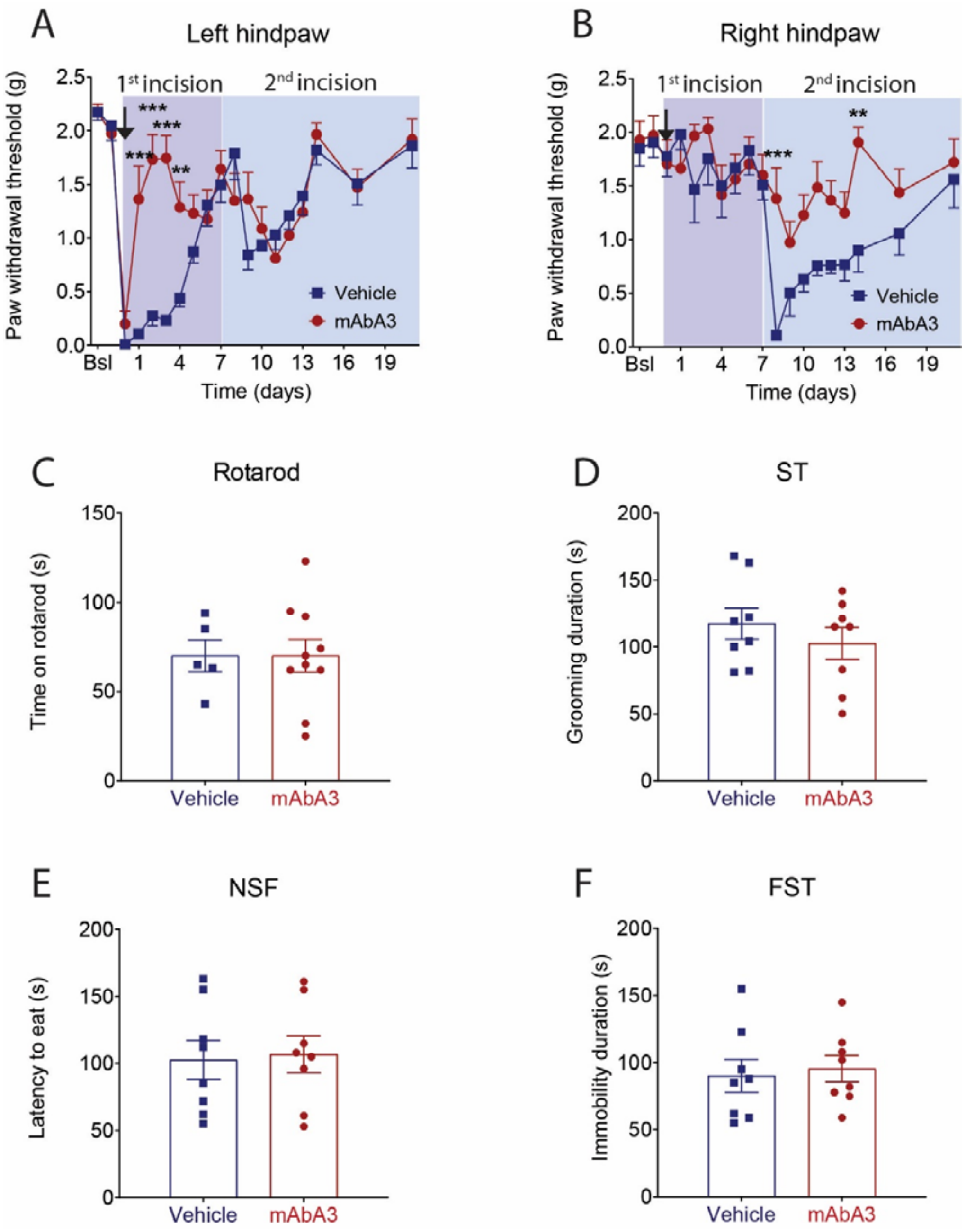


**Supplementary figure S9:** Single mAbA3 injection alleviates post-operative recovery but is not sufficient to prevent related anxiodepressive-like disorders. **(A-B)** Left and right hindpaw mechanical hypersensitivity after incisions in Vehicle or mAbA3 treated (arrow = 200µg/animal, intraperitoneal) mice as measured by the Von Frey test. **(C)** Single mAbA3 injection fails to affect motor coordination on rotarod, (**D**) grooming duration in splash test (ST), (**E**) latency to eat in novelty suppressed feeding test (NSF) and (**F**) immobility duration in forced swim test (FST) compared to control mice. All the values are means ± s.e.m. (n=8). Two-way ANOVA and Holm Sidak's test (**A-B**), Student's t-test (**C-E**); $^{**}P<0.01$; $^{***}P<0.001$ vs. Vehicle.

| Antigen | Specie | Dilution | Reference |
|---|---|---|---|
| **NeuN** | Guinea pig | 1:2000 | Millipore ABN90 |
| **ATF3** | Rabbit | 1:500 | Santa Cruz 188 |
| **CSF1** | Goat | 1:500 | R&D AF416 |
| **IBA1** | Rabbit | 1:500 | Wako |
| **Brdu** | Rat | 1:500 | Abcam ab6326 |
| **CGRP** | Goat | 1:300 | Abcam ab36001 |
| **GFAP** | Chicken | 1:1000 | Abcam ab4674 |

**Table S1:** Antibodies list for immunohistochemistry

| Gene | Forward primers | Reverse primers |
|---|---|---|
| ***Atf3*** | ACAACAGACCCCTGGAGATG | CCTTCAGCTCAGCATTCACA |
| ***Sprr1a*** | CCAGCAGAAGACAAAGCAGA | GGGCAATGTTAAGAGGCTCA |
| ***Cd11b*** | ACATGTGAGCCCCATAAAGC | AATGACCCCTGCTCTGTCTG |
| ***Iba1*** | GGATCAACAAGCAATTCCTCGA | AGCCACTGGACACCTCTCTA |
| ***Csf1*** | TGCTAAGTGCTCTAGCCGAG | CCCCCAACAGTCAGCAAGAC |
| ***Csf1r*** | ACACGCACGGCCACCATGAA | GCATGGACCGTGAGGATGAGGC |
| ***Gfap*** | GCCACCAGTAACATGCAAGA | GCTCTAGGGACTCGTTCGTG |
| ***Flt3*** | ATCCCCAGAAGACCTCCAGT | CTGGGTCTCTGTCACGTTCA |
| ***Fl*** | TTGTGGAGCCTCTTCCTAGC | AGGTGGGAGATGTTGGTCTG |
| ***Ube2e3*** | TGCTGGGCCTAAAGGAGATA | TCCCTGACTGTTGATGTTGC |
| ***Ywaz*** | TAGGTCATCGTGGAGGGTCG | GAAGCATTGGGGATCAAGAACTT |

**Table S2:** Primers list for RT-qPCR

### Supplementary methods and materials

#### Production of human recombinant FL (rh-FLT3-L)

Recombinant FL was produced in the E. coli Rosetta (DE3) strain (Novagen) in our laboratory using the pET15b-rhFL plasmid according to the protocol described [1] with some minor modifications. The rh-FL was checked for endotoxin content using the Pyrogen Recombinant Factor C endotoxin detection assay from LONZA (Walkersville MD, USA) and was found free of endotoxins.

#### Anti-FLT3 antibody development and production

Three anti-FLT3 human and murine cross-reacting scFv antibodies were selected by phage display from the human scFv synthetic library HuscI [2, 3] after sequential panning against recombinant human-FLT3-hFc and murine FLT3-hFc (R&D systems). The human scFv were reformatted as a chimeric human/murine IgG2a with Fc N297A mutation to block FcgR binding and thus antibody-mediated immune cells and complement recruitment. Recombinant mAb was produced using HEK-293T cells after transfection with polyethylenimine PEI (Polyscience) and purified using protein A chromatography. Antibody was eluted at acidic pH (glycine.HCl pH 2.7), and the solution immediately neutralized with Tris.HCl buffer, pH 9.0. Antibody was concentrated and dialyzed against PBS using an ultrafiltration centrifugal device with a cut-off of 50 kDa, sterilized by 0.2 µm filtration and stored at 4 °C. A test for the presence of endotoxins was performed and was lower than 0.25 EU/mg.

#### Behavioral testing

*Mechanical nociception assay*: Tactile withdrawal threshold was determined in response to probing of the hindpaw with eight calibrated von Frey filaments (Stoeling, Wood Dale, IL, USA in logarithmically spaced increments ranging from 0.04 to 8 g. Filaments were applied perpendicularly to the plantar surface of the paw. The 50% paw withdrawal threshold was determined in grams by the Dixon nonparametric test [4]. The protocol was repeated until four changes in behavior occurred.

*Heat test:* A radiant heat source (plantar test Apparatus, IITC Life Science, Woodland Hills, USA) was focused onto the plantar surface of the paw. The paw withdrawal latency was recorded. A maximal cut-off of 20 s was used to prevent tissue damage.

*Anxiodepression testing*: Splash test (ST), Novelty suppressed feeding test (NSF) and Forced swim test (FST) were conducted as previously described [5–7].

*Rotarod test*. The speed was set at 10 rpm for 60 s and subsequently accelerated to 80 rpm over 5 min. The time taken for mice to fall after the beginning of the acceleration was recorded.

*Conditioned place preference* (CPP). Pain-induced tonic aversive state can be unmasked by the administration of non-rewarding and rapidly acting analgesic drugs such as clonidine. All experiments were conducted by using the single trial CPP protocol as described previously for rodents [8]. The apparatus (bioseb) consists of 2 chambers (size 20 cm x 18cm x 25 cm) distinguished by the texture of the floor and by the wall patterns connected to each other by a central chamber (size 20 cm x 7 cm x 25 cm). First, animals went through a 3-day pre-conditioning period with full access to all chambers for 20 minutes. Mice underwent surgery (SI or DI) the first day of this period. At this step, mice with a spontaneous preference up to 75% were removed from the experiment. Day 4 is the conditioning day. Briefly, mice received saline intrathecal injection (5µl) and were restricted for 15 minutes to one chamber. 4 hours later, mice received clonidine intrathecal injection (1µg/5µl) and were restricted for 15 minutes to the opposite chamber. On the test day (d5), 20 h after the afternoon pairing and 4 days after surgery, mice were placed in the middle chamber of the CPP box with all doors open so animals could have free access to all chambers. The time spent in each chamber was recorded for 20 min for analysis of chamber preference.

**Immunohistochemistry**

Mice were transcardially perfused with PBS followed by 4% formaldehyde for all experiments. DRG (L4-L6), spinal cord (lumbar segment), and brains were collected and post-fixed in 4% paraformaldehyde between 10 min to 12 hours depending on the antibody and tissue before being cryoprotected in 30% sucrose in PBS. Tissues were then frozen in O.C.T (Sakura Finetek). Sections (DRG : 12 µm ; Spinal cord: 14µm) were prepared using the Cryostat Leica CM2800E. Brain sections were prepared using a Vibratome. For immunostaining, frozen sections were blocked and permeabilized in Ca2+/Mg2+-free PBS (PBS 1× −/−) containing 10% donkey serum and 0.1% or 0.3% Triton x-100 during 30 min. The sections were then incubated with primary antibodies, at 4 °C overnight in PBS 1× −/− containing 1% donkey serum and 0.1%, 0.01, or 0.03% Triton x-100. After extensive wash in PBS 1× −/− (3 times for 10 min minimum each) sections were incubated with appropriate secondary antibody conjugated to AlexaFluor and Hoechst (Sigma 1 µg/ml), in the same buffer as the primary antibodies, at room temperature for 1 h and then washing (three times for 15 min each) before mounting with Mowiol.

Immunostainings were performed with the following antibodies (see table 1): ATF3 (Santa Cruz), CSF1 (R&D), Iba1 (Wako), GFAP (abcam), CGRP (abcam), NeuN (Millipore), BrdU (Abcam), and fluorophore coupled secondary antibodies (Alexa Fluor 488, 555, 594, 647) (Invitrogen). Images were collected with a Carl Zeiss LSM 700 microscope or a Zeiss Axioscan slides scanner and were processed with Fiji/ImageJ (NIH). Corresponding images (e.g. ipsilateral vs. contralateral; FL vs vehicle; wt vs. mutant) were processed in an identical manner. Each experiment was performed in at least 3 animals. For proliferation assays, BrdU (Sigma) was injected intraperitoneally into adult mice (2 mg/animal), 24 hr before sacrifice. For BrdU detection, slides were incubated in 10 mM sodium citrate buffer (pH 6.0) at 95°C for 15 min, then at room temperature (RT) for 15 min and subjected to immunostaining.
